# Stability and diversity of interactions in complex microbiomes

**DOI:** 10.1101/2025.09.24.678377

**Authors:** Jorge Calle-Espinosa, Jaime Iranzo

**Affiliations:** Centro de Biotecnología y Genómica de Plantas, Universidad Politécnica de Madrid (UPM)-Instituto Nacional de Investigación y Tecnología Agraria y Alimentaria (INIA-CSIC), Madrid, Spain; Centro de Astrobiología (CAB) CSIC-INTA, Torrejón de Ardoz, Madrid, Spain; Institute for Biocomputation and Physics of Complex Systems (BIFI), University of Zaragoza, Zaragoza, Spain

**Keywords:** microbial community, interaction network, community dynamics, cross-feeding, microbiome engineering, high-order interaction, mutualism, competition, biotechnology

## Abstract

Complex microbial communities abound in nature, yet designing robust and stable microbial consortia for use in biotechnology and One Health applications remains a major challenge. Inspired by a large body of work on community ecology, we used mathematical models to investigate what makes a microbiome stable. Our approach integrates three factors that characterize many free-living and host-associated microbial communities but have seldom been jointly considered: sustained growth, high-order interactions, and a broad diversity of interaction profiles. We derived a simple expression that relates community stability to basic statistical properties of the underlying interaction networks. Analytical and numerical results show that synergistic high-order interactions, such as those derived from complex auxotrophies, play a critical role in stabilizing species-rich microbial communities with highly diverse interaction profiles. In contrast, in the absence of high-order interactions, viability is restricted to communities with very sparse or fully mutualistic interaction networks. Our findings have the potential to inform microorganism-based interventions in areas in which sustained growth and long-term stability are desirable properties. We propose that incorporating high-order mutualism into engineered microbial consortia could improve their robustness by buffering design uncertainty about the underlying interaction networks.

## Introduction

Microorganisms are ubiquitous in nature. They play central roles in the biosphere by fixing carbon and nitrogen, facilitating nutrient cycling and establishing mutualistic relationships with plants and animals [1–4]. Recent advances in environmental meta’omics and culture techniques have shed light on the complex structure of many microbial communities and the tight coupling of microbiome dynamics with host-level and ecosystem-level functions [5–7]. Thus, transitions between healthy and perturbed states of macroscopic systems are often associated with the disruption of resident microbial communities, with eutrophication and gut dysbiosis as paradigmatic examples [8–10]. Along with these findings, microbiome engineering has emerged as a promising strategy to improve human health [11], increase crop productivity [12], prevent biodiversity loss [13, 14], and mitigate climate change [15]. Microbiome engineering aims to control and enhance the function of natural microbial communities by adding microorganisms or modifying those that are already present. To produce sustained benefits, engineered microorganisms must integrate into the community and persist in the long term. The stable integration of new microorganisms in natural and synthetic microbiomes is a major unsolved challenge in microbiome engineering, which has limited its scope to short-term interventions [16].

The challenge of designing persistent microbial consortia is linked to a central problem in community ecology: identifying the general principles that determine the stability of biological communities and their sensitivity to the introduction of new species. From a theoretical perspective, the conceptualization of the paradox of biodiversity, which predicts a complexity threshold beyond which ecosystems collapse, was a milestone in the field [17, 18]. Since then, much work on theoretical ecology has dealt with explaining why complex communities exist [19]. Recently, the study of microbiomes has benefited from models and conceptual frameworks originally developed for community ecology [20, 21]. These have been applied, for example, to explaining trends in microbial diversity [22–24] and understanding connections between metabolism and community dynamics [25, 26]. Taking a further step in that direction, we adapted models from community ecology to understand how biotic interactions determine the stability of complex microbiomes and, consequently, which ecological structures are more likely to result in stable microbial communities. To do so, we considered three elements that are central to microbial communities but have seldom been jointly considered by previous works: sustained growth, high-order interactions, and interaction diversity.

Sustained growth characterizes most engineered microbial communities used in food production, wastewater treatment, and biotechnological applications. Sustained microbial growth can also occur in natural environments, like the gut or aquatic systems, that are maintained in a non-equilibrium state by a more or less continuous supply of nutrients and removal of microbial biomass through outflow or sedimentation. Importantly, not all modeling frameworks are equally suited to accommodate scenarios of sustained population growth. For example, generalized Lotka-Volterra equations, which are widely used to infer microbial interaction networks from time series [27, 28], impose by design a zero-growth constraint on the average abundances at equilibrium. Because of that, feasibility (the existence of steady states with survival of all the species) and stability in Lotka-Volterra models are strongly associated with conditions that prevent unbounded proliferation, such as saturating growth functions [29], weak interactions [30–34], widespread competition [22], or diminishing returns for mutualism [35–37]. In contrast, generalized replicator equations, originally developed in the context of evolutionary game theory, naturally accommodate scenarios in which the population grows while the species’ relative abundances remain constant or fluctuate around an equilibrium composition [38, 39].

A second aspect that characterizes microbial communities is the relevance of high-order interactions, that is, interactions that involve three or more species. These emerge, for example, in scenarios in which metabolic complementation between two species requires the presence of a third species [40–42]. High-order interactions can also appear in the context of microbial warfare if sensitivity to antibiotics produced by a competitor varies depending on the presence of additional biotic factors, such as bacteriocins or antibiotic-degrading enzymes released by other species [43, 44]. Though often neglected by ecological studies, high-order interactions are widespread in natural and engineered microbial consortia [45], as revealed by frequent inconsistencies between the dynamics of the full community and isolated pairs of species [46–49]. Mathematical models suggest that high-order interactions have the potential to shape ecosystem diversity and promote coexistence [50–53]. Although the experimental evidence is still limited, available data from reconstituted microbiomes supports the idea that multispecies interactions positively contribute to the stability of microbial consortia [48, 49, 54].

Finally, interaction diversity (that is, the relative prevalence of mutualism, competition, and exploitation) has received much attention in the context of ecological stability [30, 33, 35, 55, 56], but rarely in combination with sustained growth or high-order interactions. Likewise, previous research on the effect of high-order interactions has typically focused on a rather limited range of scenarios, described by zero-sum competitive networks or zero-mean random matrices. Although the relative weight of mutualism and competition in complex microbiomes is a matter of debate [57–62], it is unlikely that positive and negative interactions are balanced in general. Moreover, interventions in microbiome engineering harness a broad range of interaction effects, from facilitative cross-feeding in bioreactors to antagonistic microbial warfare in therapeutics [63]. Such diversity warrants a generalized approach to the study of microbiome stability beyond the constraints of traditional (Lotka-Volterra, balanced, or pairwise interaction) frameworks.

## Results

### Replicator model for the growth of complex microbial communities

To investigate how different types and orders of interaction affect the stability of microbial communities, we considered a community of *N* species (understood as ecological units) whose dynamics are described by a system of generalized replicator equations:

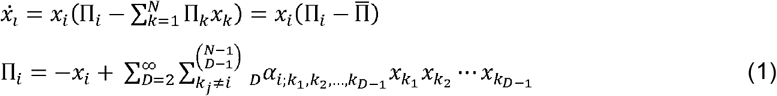

In this model, the relative abundance of a species *i*, denoted as *x*_*i*_, increases or decreases at a rate that is proportional to its frequency and the difference between its fitness (that is, its effective growth rate, Π) and the mean fitness of the community 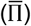. The fitness of each species depends on the composition of the community through a network of positive and negative interactions. Such interactions, represented by the _*D*_*α*_*i*;k_ factors in eqn. 1, are not necessarily symmetric and can involve an arbitrary number of species (*D*). The model includes a self-inhibition term to reflect the assumption that species are self-limiting at high concentration, as expected if members of the same species share the same niche. Self-inhibition guarantees stability in the absence of interactions. Without loss of generality, all interactions are rescaled with respect to the strength of the self-inhibition term, which takes a value of one. This is equivalent to measuring the interaction strengths in terms of the self-inhibition term. Following previous studies [22, 33, 35, 64, 65], to single-out the effect of interactions, we assumed that the basal growth rate in the absence of interactions is the same for all the species (with that assumption, the dynamics become invariant with respect to the basal growth rate, so we set it to zero for simplicity).

Cross-species interactions can be characterized by the benefit or disadvantage that they entail for each interactor. Thus, pairwise (*D*= 2) interactions can be mutualistic (+/+), competitive (-/-), exploitative (+/-), commensal (+/0), or amensal (0/-). In a similar fashion, higher order interactions (*D*> 2) can be classified according to the number of species that benefit, lose, and remain unaffected by the interaction (Fig. 1). For simplicity, we assumed that interaction networks are random and independent across orders, and the interaction strength takes a fixed value ± *ϕ*_*D*_ for interactions involving *D* species. Moreover, we focused on interactions that affect all the participant species (i.e., high-order analogs of mutualism, competition, and exploitation), although analytical expressions for high-order commensalism and amensalism are included in the Supplementary Information. Under these assumptions, the statistical properties of any interaction network can be described by three parameters: the positivity (*P*_*D*_), the symmetry (*S*_*v*_), and the connectivity ( *C*_*D*_), where the subindex *D* specifies the order of the interaction. (An additional parameter would be required to describe generalized forms of mensalism, see SI, section 4.1.) The positivity, defined as the fraction of ‘+’ signs in the network, determines the mean benefit that a species gets from ecological interactions. The symmetry is the probability that two interacting species are affected in the same way, either positively or negatively. Therefore, the more exploitative the network, the lower the symmetry. Finally, the connectivity is the fraction of all possible interactions that are not null. Positivity, symmetry, and connectivity take values between 0 and 1, but not all combinations are feasible. For example, a network with full positivity (*P*_*D*_ = 1) only comprises mutualistic interactions, which implies full symmetry (*S*_*D*_ = 1). In general, the more extreme the positivity and the symmetry, the less diverse the spectrum of ecological interactions.

**Figure 1.**
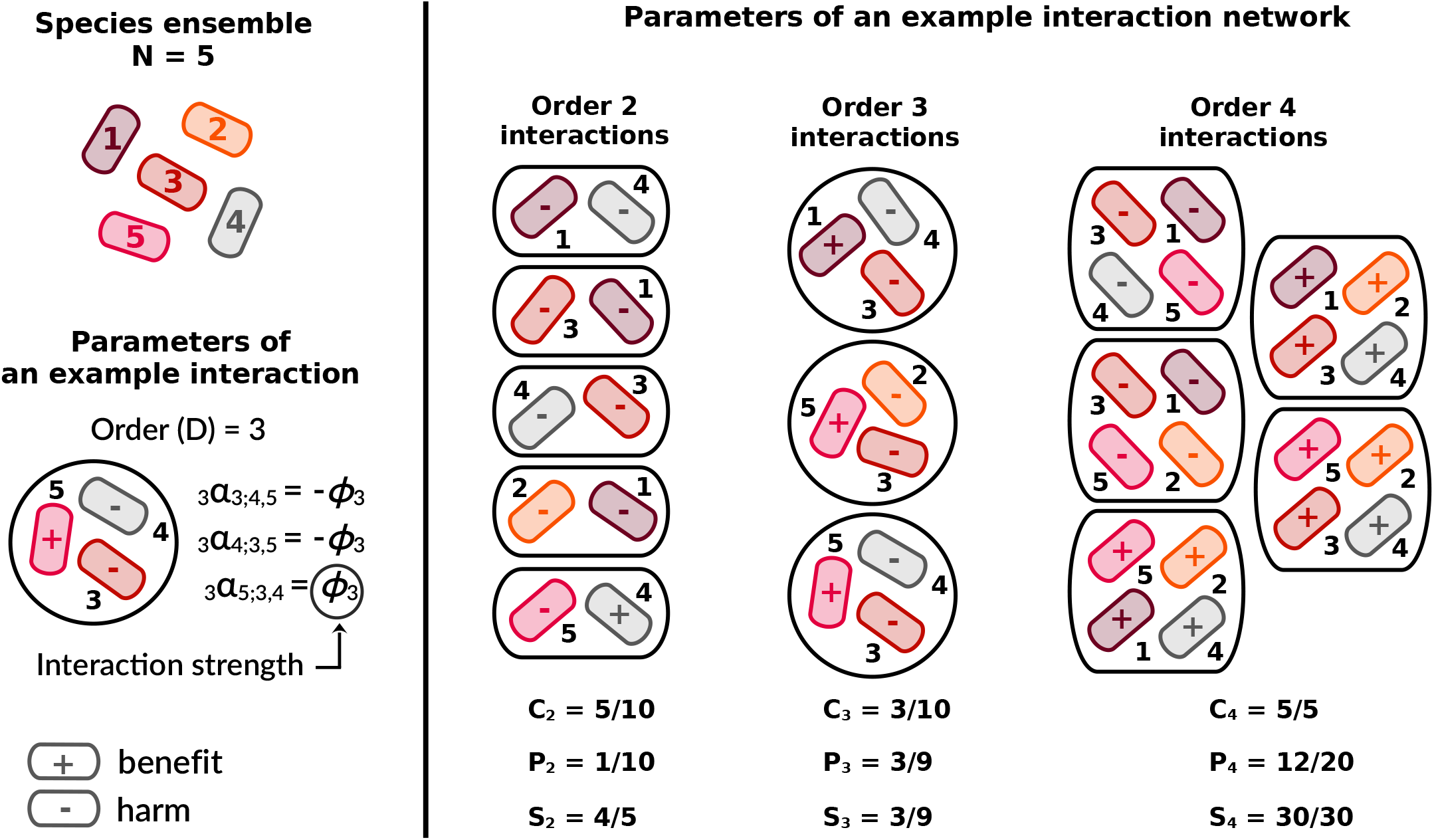
Characterization of high-order interaction networks. An example network with *N* = 5 species and interactions of order 2, 3, and 4. Each species is numbered and represented as a rounded colored rectangle. Interactions are represented as groups of species enclosed by a black line. Given an interaction, the + and − symbols indicate if a species is positively or negatively affected by that interaction. Left, *Parameters of an example interaction*: An order *D* interaction involving species *i,k*_1_, *k*_2_,…, *k*_D − 1_ (species nos. 3, 4, and 5 in this example, order *D* = 3) is completely described by a set of *D* constants 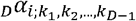 that capture how each species is affected by that interaction. In this study, we assume that the magnitude of these constants (the interaction strength) is fixed for each order (±*ϕ*_*D*_). Main panel, *Parameters of an example interaction network*: In this example, connectivity ( *C*_*D*_), positivity (*P*_*D*_), and symmetry (*S*_*D*_) are different for each interaction order. Note that positivity and symmetry are not completely independent: extreme positivity (close to 0 or 1) implies high symmetry; although the opposite is not true. This is illustrated by order-2 and order-4 subnetworks, respectively. Note also that, at interaction orders ≥3, the symmetry is always greater than zero.

### Stability of microbial communities at different interaction orders

Ecological stability is a complex and multifaceted concept [66]. Here we focus on local asymptotic stability, defined as the ability of a system to return to its original equilibrium state after (infinitesimally) small perturbations. Local asymptotic stability represents a necessary condition that must be held so that other, more nuanced components of stability (such as resilience and tolerance) can be evaluated. By applying linear stability analysis (SI, section 3.1), we obtained the following approximate condition for the asymptotic stability of a large community:

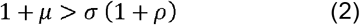

where

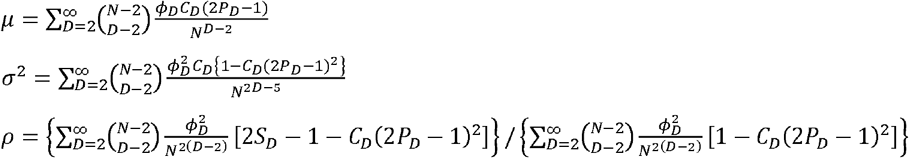

The stability condition includes three terms, *μ, σ*, and *ρ*. The first two represent the mean (*μ*) and standard deviation (*σ*) of the fitness contribution that a species receives from all other species, considering all possible interaction orders. The third term (*ρ*) represents the correlation of reciprocal fitness contributions calculated over all pairs of species. That is, *ρ*= 1 if both species of all pairs affect each other in the same way, *ρ*= −1 if they affect each other by the same amount but with opposite sign, and *ρ*= 0 if fitness contributions are uncorrelated. It is clear from eqn. 2 that stability is associated with large means, small deviances, and negative correlation of fitness effects.

To verify the validity of eqn. 2, we numerically integrated the replicator equations under a wide range of community sizes and interaction profiles (SI, section 3.2 and Fig. S1). The numerical results confirm that eqn. 2 provides an accurate, though slightly conservative condition for the stability of a community. Violations of this criterion are associated with small community sizes (low *N*) and parameter regions where the predicted range of stable networks is extremely narrow. In those conditions, stable communities can sometimes be found even if eqn. 2 does not hold.

If only one order of interaction is considered, it follows from eqn. 2 that increasing the interaction strength (*ϕ*_*D*_) always works against stability. In consequence, the maximum value of *ϕ*_*D*_ that is compatible with stability 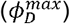 is a good measure of the viability of an interaction network. In general, communities enriched in exploitative (low symmetry) and cooperative (high positivity) interactions are the ones that achieve the highest viability (Fig. 2A). In contrast, competitive communities are typically unstable, unless the interaction network is very sparse. Such low stability results from the mean term of eqn. 2 becoming negative once cross-species interactions overpower self-inhibition. Network connectivity has a complex impact on stability, which is explained by its dual effect on the mean and deviance terms of eqn. 2. On one hand, connectivity contributes to the absolute value of the mean effect size, reinforcing the stabilizing or destabilizing effects of cooperation and competition, respectively. On the other hand, high connectivity promotes stability by reducing the deviance term in purely competitive or cooperative networks, but it has the opposite effect in balanced or exploitative networks. All in all, low connectivity generally favors stability, with a stronger effect in competitive networks, whereas high connectivity promotes stability in highly cooperative networks.

**Figure 2.**
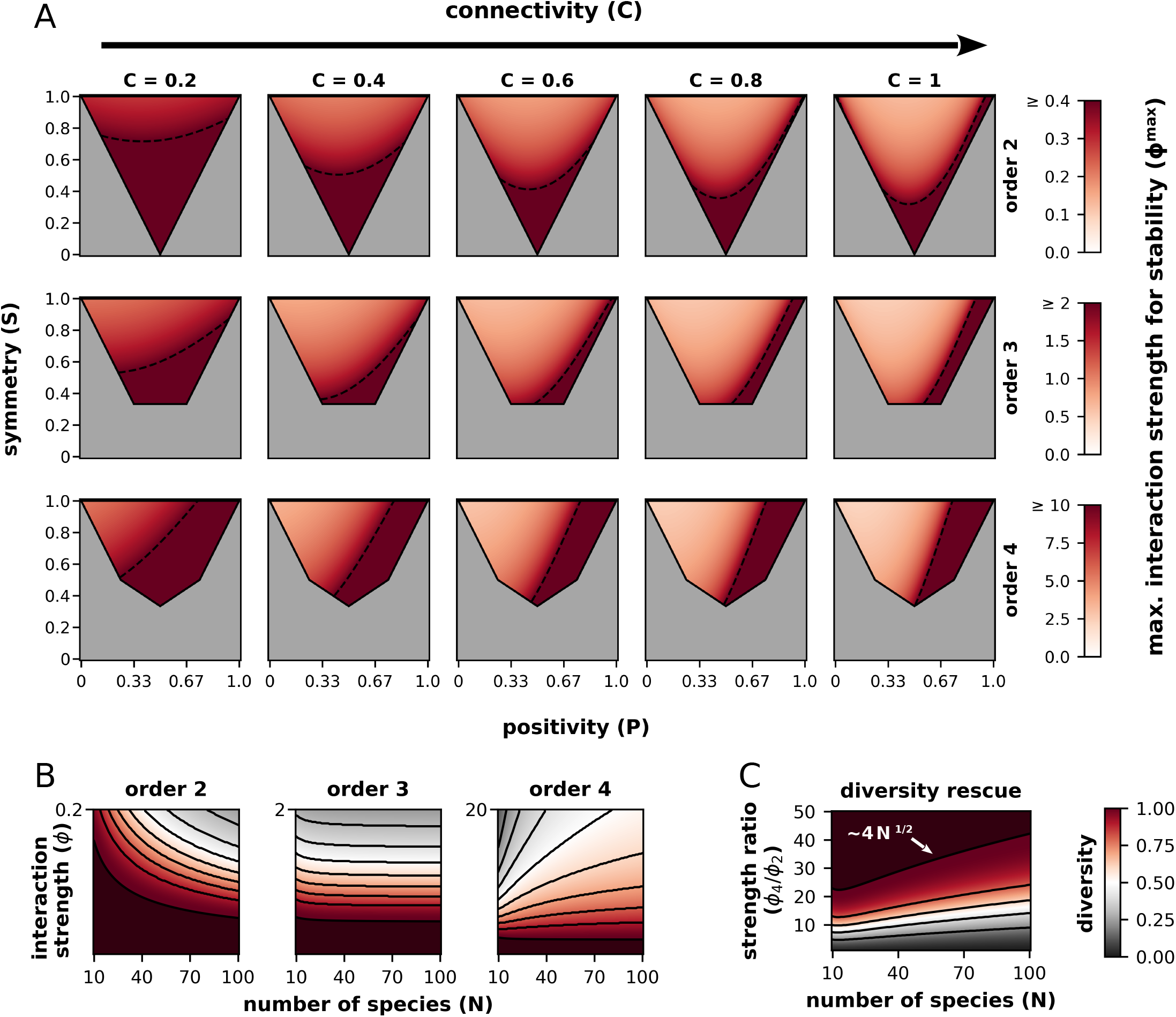
Stability and diversity of interactions in complex communities. (A) Maximum interaction strength that is compatible with stability 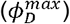 at different interaction orders and a wide range of connectivities, positivities, and symmetries (*N* = 15 species). Dashed lines correspond to 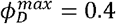 (order 2), 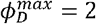 (order 3), and 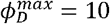 (order 3). Combinations of positivity and symmetry that cannot be realized (by definition) are displayed in gray. (B) Diversity of interaction profiles, measured as the fraction of all possible interaction profiles (realizable combinations of connectivity, positivity and symmetry) that are stable. The diversity of interaction profiles is displayed for single-order networks and different values of the species richness (*N*) and interaction strengths (*ϕ*_*D*_). Black lines represent level curves associated with different diversity values. Decreasing level curves (order 2) indicate that as the species richness increases, the range of interaction profiles that are compatible with stability shrinks; increasing level curves (order 4) denote the opposite trend; horizontal level curves (order 3) imply insensitivity with respect to species richness. (C) Rescue of unstable order-2 networks by adding high-order interactions. The plot shows the fraction of interaction profiles of order 2 (with *ϕ*_2_ = 1) that become stable after superimposing a fully connected, completely mutualistic order-4 interaction network ( *C*_4_ = 1, *P*_4_ = 1, *S*_4_ = 1) as a function of the species richness (N) and the ratio of order-4 vs. order-2 interaction strengths (*ϕ*_4_/ *ϕ*_2_). Black lines represent level curves associated with different diversity values.

### Interaction order critically determines diversity-vs-stability trade-offs

A remarkable outcome of this analysis is that the association between species richness and viability depends on the order of interaction. Of the three factors that control stability, the deviance is the only one that strongly depends on the number of species, scaling as 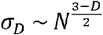 for order *D* interactions (SI, section 3.1). This implies that, for pairwise interactions, increasing the number of species destabilizes the community, whereas the opposite holds for interaction orders ≥4. (Species richness does not significantly affect stability for order 3 interactions.) The critical reversal of the diversity-vs-stability trade-off at high interaction orders does not only allow for higher species richness (as previously reported in [51]), but also for more diverse interaction profiles. To illustrate that point, we calculated, for different values of species richness and interaction strength, the fraction of all possible interaction profiles, in terms of connectivity, positivity and symmetry, that are stable. (Such a fraction represents an alpha-diversity for interaction profiles; thus, we refer to it as “interaction diversity”.) As shown in Fig. 2B, given a fixed interaction strength, the interaction diversity of stable communities decreases, remains stable, and increases with species richness in 2nd-order, 3rd-order, and 4th-order interaction networks, respectively. Conversely, the maximum interaction strength compatible with a given interaction diversity (level curves in Fig. 2B) decreases, remains stable, and increases with species richness in 2nd-order, 3rd-order, and 4th-order networks.

In summary, in pairwise interaction networks, stability is usually associated with low species richness, sparsity, or homogeneity of interactions (fully exploitative or mutualistic). In contrast, high-order interaction networks (especially those with interaction orders ≥4) can accommodate much higher heterogeneity while remaining stable. By taking the limit N → ∞, the stability condition for order ≥ 4 becomes

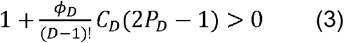

Accordingly, high-order networks in which cooperation is dominant 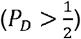 are always stable if the number of species is sufficiently large, regardless of their connectivity. In contrast, the stability of large, high-order competitive networks is not always guaranteed, although it can be reasonably expected unless interactions are too strong.

### Diversity rescue driven by high-order interactions

The trends described so far apply if all interactions belong to a single order, but what if a mixture of interactions of different orders is considered? By applying similar scaling arguments as those presented above, it follows that, as the number of species increases, the relative contribution of high-order interactions to the deviance and correlation terms of eqn. 2 becomes less and less relevant. As a result, in species-rich communities, those terms are determined by the lowest non-negligible interaction order. Such lowest order typically corresponds to pairwise interactions, which work against stability as the number of species increases. In contrast, all interaction orders contribute to the mean term, promoting or hindering stability (depending on the overall sign) in a way that is almost independent of species richness. In practice, this implies that high-order cooperative interactions, if sufficiently strong or dense, can partially compensate for the destabilizing effect of pairwise interactions (Fig. 2C). The feasibility of such high-order-driven “diversity rescue” can be assessed by the ratio of high-order vs. order-2 interaction strengths required to stabilize a community of *N* species. Scaling analysis shows that such ratio grows as the square root of the number of species. Thus, a *k*-fold increase in the strength (or connectivity) of high-order interactions is sufficient to support a *k*^2^-fold increase in the species richness.

### Feasibility of stable communities

The stability condition of eqn. 2 was derived under the customary assumption that, given an interaction network, the model has a solution in which all species are present. If such a solution exists, then the stability condition guarantees that the community will be robust with respect to small perturbations. However, not all interaction networks are compatible with full-survival solutions. Obtaining a general condition that ensures persistence of all the species in the community is technically challenging. Nevertheless, by applying the Kantorovich theorem and some reasonable approximations (SI, section 2), we derived a conservative expression that proves that full survival in species-rich communities is almost always guaranteed if high-order (*D* ≥ 4) interactions are dominant. Moreover, in those conditions, the composition of the stationary state becomes more homogeneous the higher the number of species.

To investigate the general case, including non-negligible pairwise and 3rd-order interactions, we resorted to simulations. For each set of parameters, we simulated 100 interaction networks and defined the critical interaction strength at order 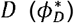 as the strength at which half of the networks produced a stable community with persistence of all the species. This value is analogous to the maximum strength 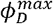 discussed above, in the sense that it serves as a proxy for the viability (now understood as feasibility and stability) of a microbial community.

Simulations confirmed that, after accounting for feasibility, the viability still follows the trends predicted by the stability criterion (Fig. 3), although with some deviations at low interaction orders. The most remarkable exception corresponds to dense exploitative networks, which fulfil the stability condition but are not feasible (Fig. 3A). Moreover, at low interaction orders, the viability of random communities is systematically lower than predicted based on stability arguments alone. Despite these discordances, the overall effect of species richness on viability at different interaction orders (Fig. 3B) and the potential of high-order interactions to rescue diversity (Fig. 3C) remain essentially unchanged.

**Figure 3.**
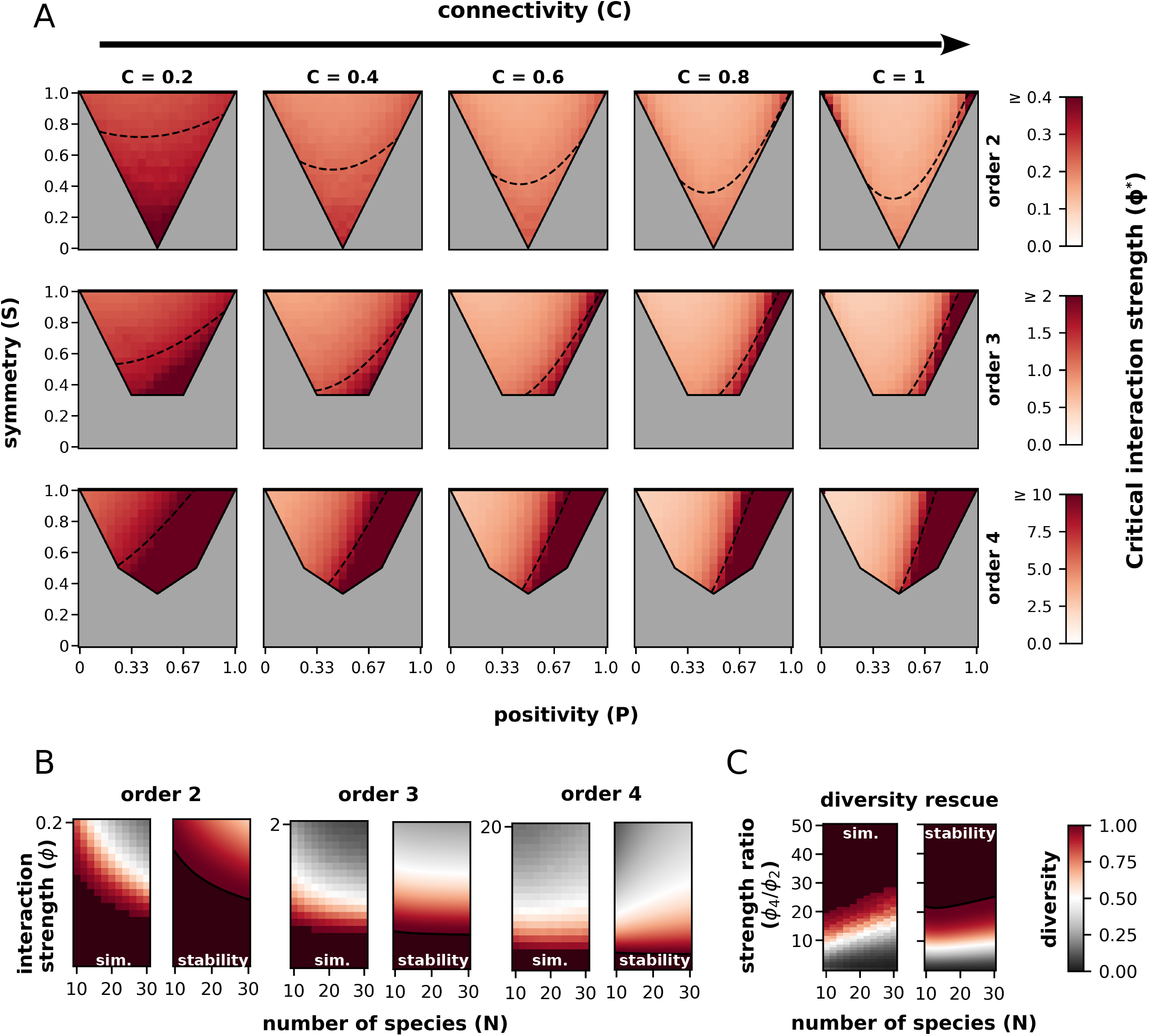
Simulation-based study of community viability and interaction diversity. (A) Critical interaction strength 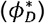 in single-order networks as a function of the interaction profile (*N* = 15 species). The critical interaction strength is defined as the highest value of the interaction strength at which at least 50% of the simulated networks reach a stable stationary state. Dashed lines correspond to analytical estimates of 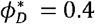 (order 2), 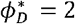 (order 3), and 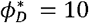 (order 3) based on eqn. 2 and are identical to those in Fig. 2. Combinations of positivity and symmetry that cannot be realized (by definition) are displayed in gray. (B) Diversity of interaction profiles, measured as the fraction of all realizable combinations of connectivity, positivity, and symmetry that are stable. The diversity of interaction profiles is displayed for single-order networks and different values of the species richness (*N*) and interaction strengths (*ϕ*_*D*_). For each interaction order, the left plot shows simulation results, and the right plot shows analytical estimates based on eqn. 2. (C) Rescue of unstable order-2 networks by adding high-order interactions. Both plots show the fraction of interaction profiles of order 2 (with *ϕ*_2_ = 1) that become viable after superimposing a fully connected, completely mutualistic order-4 interaction network ( *C*_4_ = 1, *P*_4_ = 1, *S*_4_ = 1) as a function of the species richness (N) and the ratio of order-4 vs order-2 interaction strengths (*ϕ*_4_/*ϕ*_2_). Left and right plots are based on numerical simulations and analytical estimates (eqn. 2), respectively.

## Discussion

Sustained growth, functional diversity, and high-order interactions characterize many natural and engineered microbiomes. By combining mathematical analysis and numerical simulations, we investigated how those three factors determine the feasibility and asymptotic stability of microbial communities. In the absence of high-order interactions, viable communities are limited to two types of structures: fully synergistic consortia and weakly or non-interacting systems. This result is consistent with the outcome of more detailed consumer-resource models [55] and empirical estimates of pairwise interactions in some natural communities [67–70]. High-order interactions substantially modify that picture, facilitating the emergence of species-rich microbiomes that can display a broad spectrum of interactions. This finding, that expands on previous results [51], predicts a reversal of the complexity paradox at high interaction orders and an amelioration (though not full reversal) of it in complex scenarios with mixed interaction orders. In consequence, high-order interactions can stabilize microbiomes that would otherwise be unstable, especially if such interactions are strong and involve high-order analogs of mutualism. Such a stabilizing effect has been experimentally observed in the zebrafish gut microbiome, where high-order interactions promote coexistence by dampening cross-species competition [48].

From a mathematical perspective, the reversal of the complexity paradox at high interaction orders results from a fundamental statistical property: given random interaction terms of order >3 and considering their net contributions to each species’ fitness, both the variance of such contributions and their correlation across pairs of species tend to zero as the number of species grows. This homogenizing effect is reminiscent of previous theoretical arguments whereby highly dimensional ecological niches [71] and emergent neutrality [72] support biodiversity in species-rich ecosystems.

Unlike in Lotka-Volterra models, interaction terms in replicator equations depend on the relative abundances of species rather than their densities. Classical applications of replicator equations include continuous flow reactors and spatially extended systems, such as the gut, in which the flux of biomass to adjacent regions keeps the local density constant [73]. A notable property of replicator equations is that the right-hand side can be multiplied by any arbitrary function without altering the stationary states or their stability (such transformation amounts to dynamically changing the timescale). In consequence, some forms of global regulation, such as a total biomass-dependent growth rate, can be introduced in the model without affecting the conclusions. Though not completely equivalent, systems subject to serial dilution with intermittent nutrient supply are also well approximated by replicator equations [74]. More generally, in scenarios in which cell density varies with time, relative abundances can determine population-level interaction strengths if local cell densities are high and interactions are effectively short-range, for example due to fast consumption of resources involved in cross-feeding.

The replicator equations at order 2 are structurally analogous to the compositional Lotka-Volterra model, which was recently proposed to infer ecological interactions from compositional time series [67]. In fact, both approaches converge if the average growth rate is zero, which is the scenario in which the compositional Lotka-Volterra model was originally formalized. At higher orders, or in populations with nonzero growth rates, the mathematical equivalence between replicator and Lotka-Volterra equations (both compositional and traditional) involves complex transformations of the parameters [75]. Such nontrivial correspondence may explain why some of the conclusions of this work are not observed in high-order Lotka-Volterra systems [37].

An interesting possibility that was not directly addressed in this work is that interaction terms at different orders scale with the volume or cell density of the system. For example, it has been shown through micro-scale bioprinting of microbial colonies that metabolite sharing is distance-dependent and the presence of competitors along the path of metabolite diffusion can curtail cross-feeding [76]. Because high-order interactions involve a higher number of species and extracellular compounds, it can be hypothesized that they decay faster with distance and dilution than pairwise interactions. That would result in a scenario characterized by short-range (or high-density) stability, provided that high-order interactions are sufficiently strong, but long-range (or low-density) instability once pairwise interactions become dominant. Such multi-scale behavior could be relevant to understanding the dynamics of spatially heterogeneous microbiomes, such as kefir, that include matrix-bound (grains) and liquid phases [77].

The stabilizing effect of high-order mutualism has relevant implications for the design of microbial consortia. Incomplete knowledge of ecological interactions, environmental perturbations, and evolutionary change are sources of uncertainty that can compromise the success of microbiome-based interventions. By expanding the spectrum of viable interaction networks, multispecies interdependencies could buffer such uncertainty and reduce the risk of collapse in engineered communities. Accordingly, synthetic microbiomes encompassing networks of auxotrophies that require complementation from more than two producer species may prove substantially more robust than simpler designs based on pairwise interactions. In light of these findings, incorporating complex interdependencies in engineered microbiomes seems worth exploring despite the challenges.

## Methods

### Generation of random interaction networks with desired properties

Given an interaction order *D*, there are *D*+ 1 possible classes of interactions (excluding generalized forms of mensalism) that are univocally characterized by the number of species that benefit from the interaction. To build a *D*-th order interaction network of *N* species with positivity *P*_*D*_, symmetry *S*_*D*_, and connectivity *C*_*D*_, we first obtained the fraction of interactions of each class by solving the following optimization problem:

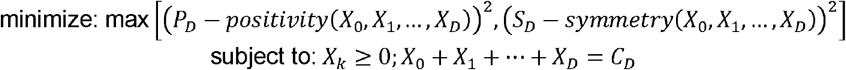

where the variable *X*_*k*_ indicates the fraction of non-null interactions where *k* species are benefited, and the “positivity” and “symmetry” functions are calculated according to the definition of these properties (Figure 1 and SI section 4.1l). Since we only consider compatible pairs (*P*_*D*_,*S*_*D*_), the minimum of this expression is necessarily zero. The minimization problem was solved using the *COBYLA* method implemented in the *minimize* function of the *scipy* package (version 1.8.0). Because there may be multiple compatible solutions for the same pair (*P*_*D*_, *S*_*D*_), the initial condition of the minimization problem was randomly set by extracting the values of *X*_*k*_ from a Dirichlet distribution and normalizing them to ensure that the connectivity constraint is fulfilled. To obtain the number of interactions of each class, we multiplied the fractions (*X*_0_, *X*_1_,…, *x*_*D*_) by the total number of order *D* interactions, given by the binomial coefficient 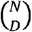. Finally, we randomly assigned each interaction to one of all the possible sets of *D* species.

In networks with multiple interaction orders, the procedure described above was separately applied to each interaction order, under the assumption that interactions are independent across orders.

### Numerical simulations of community dynamics

The replicator equations (eqn. 1) were numerically integrated using the *LSODA* method implemented in the function *solve_ivp* of the *scipy* package (version 1.8.0), with default relative and absolute tolerances and explicit Jacobian. The initial condition for all the simulations was a perturbation of the stationary state found by the Newton-Raphson method. (We used the homogeneous state, with all species in the same proportion, as the starting point for the algorithm.) The perturbation involved multiplying the composition of the stationary state by 0.999 and adding a random term drawn from a Dirichlet distribution (all parameters equal to one) multiplied by 0.001. If the Newton-Raphson method did not converge, we used the homogeneous state as the initial condition for the numerical integration.

The equations were numerically integrated until any of the following conditions were fulfilled: (i) the abundance of any species drops below 10^−6^/N (the community is unstable); (ii) the sum of the absolute values of the time-derivatives of all the species abundances is smaller than 10^−8^ (the community has a stable stationary state); (iii) the entropy (−∑x_*i*_ log(x_*i*_), where x_i_ is the proportion of species i) remains constant or bounded in its fluctuations (see details below); (iv) neither of the above criteria is fulfilled before 1 hour (real time) or 10^5^ time units (simulated time), or the sum of all the species abundances deviates from 1 by more than 0.1% (the simulation is discarded).

The entropy criterion, as previously defined [51], was applied as follows: first, the system was simulated for 1000 time units. Then, the minimum value of the entropy in the last 100 time units was compared with the minimum value at any previous time (that is, excluding the last 100 time units). The same was done with the corresponding maxima. If both ratios differed from 1 in less than 10^−8^, then the entropy was considered constant or bounded in its fluctuations and the simulation ended. Otherwise, the system was simulated for additional periods of 100 time units until it fulfilled the entropy criterion or any of the other stop criteria. Communities that fulfilled the entropy criterion were considered to have reached an attractor with all species present.

Although attractors that are not fixed-point stationary states (e.g., limit cycles or chaotic attractors) are found for some interaction networks, these are in general rare (Fig. S2). Because of that, and to facilitate comparison with analytical results (which only apply to fixed-point stationary states), we based our numerical assessment of viability on simulations that reached a stable (fixed-point) stationary state.

### Calculation of the interaction diversity

We measured the diversity of interaction profiles as the fraction of all feasible combinations of connectivity, positivity, and symmetry that fulfill the stability condition (in the analytical study, Fig. 2) or display stable species coexistence (in the simulation-based study, Fig. 3) given a particular interaction order *D*, species richness *N* and interaction strength *ϕ*_*D*_. Conceptually, this can be interpreted as a normalized alfa-diversity in the space of interaction profiles. To obtain this value, each feasible point of the space *C*_*D*_, *P*_*D*_, *S*_*D*_) was assigned a weight between 0 and 1 that represents the fraction of all possible networks compatible with those parameters that fulfil the stability criteria (note that, in the analytical study, that weight is either 0 or 1). The interaction diversity is the integral of the weight function over all possible parameter combinations, divided by the corresponding hypervolume.

The integration of the weight function in the analytical study, as well as the calculation of the total hypervolume, was performed using the *dblquad* function of the *scipy* package (version 1.8.0) with default options. A Monte-Carlo approach was implemented to calculate the interaction diversity from simulations: We uniformly sampled 10000 parameter combinations from the space (*C*_*D*_,*P*_*D*_, *S*_*D*_) and, for each combination, we obtained a random interaction network. The fraction of simulations that ended in a stable stationary state was used as an estimate of the interaction diversity.

### Estimation of the critical interaction strength 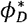

The critical interaction strength is defined (for fixed order *D*, number of species *N*, positivity *P*_*D*_, symmetry *s*_*D*_, and connectivity *C*_*D*_) as the interaction strength *ϕ*_*D*_ at which half of the simulations reach a stable stationary state. This value was obtained *via* binomial regression using the logit link; *ϕ*_*D*_ as the predictor variable, and *n*_*s*_, the number of simulations from a total of 100 that ended in a stable stationary state, as the response variable. The regression was performed using the *statsmodel* package (version 0.14.0) with default parameters. The critical interaction strength 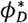 was calculated as the absolute value of the intercept divided by the coefficient of the variable *ϕ*_*D*_.

To minimize the computational cost associated with the estimation of 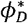, a reduced subset of *ϕ*_*D*_-values was used for the regression. To select that subset, we first applied a procedure inspired by the bisection method to find the largest *ϕ*_*D*_ at which *n*_*s*_ = 100 (*ϕ*_*low*_) and (if it exists) the smallest *ϕ*_*D*_ at which *n*_*s*_ =0 (ϕ_*upp*_). Then, we sampled 10 values of ϕ_*D*_ between *ϕ*_*low*_ and ϕ_*upp*_ to characterize the region of the *n*_*s*_ *vs ϕ*_*D*_ curve that varies the fastest. Note that, because the bisection method evaluates *n*_*s*_ for *ϕ*-values progressively closer to *ϕ*_*low*_ and ϕ_*upp*_, it is possible to further reduce the computational cost of the algorithm by reusing those *ϕ*-svalues that lie between *ϕ*_*low*_ and ϕ_*upp*_.

To obtain the (very coarse) estimate of *ϕ*_*low*_, lower (^*m*^*ϕ*_*low*_) and upper (^*M*^*ϕ*_*low*_) bounds for that value were first established. ^*m*^*ϕ*_*low*_ =0 warrants that *n*_*s*_ = 100. ^*M*^*ϕ*_*low*_ was initially set to 1.5 times the maximum *ϕ*_*D*_ compatible with stability (*ϕ*_*ref*_) according to the criteria of eqn. 2. Then, if the *n*_*s*_ associated with ^*M*^ *ϕ*_*low*_ was ≥65, a new ^*M*^ *ϕ*_*low*_ was defined by adding *ϕ*_*ref*_. This procedure was iterated until the *n*_*s*_ associated with ^*M*^ *ϕ*_*low*_ was ≤65 or until ^*M*^ *ϕ*_*low*_ ≥ 1000. In the last case, *M ϕ*_*low*_ was set to 1000. Given these boundaries, the value of *ϕ*_*low*_ was approximated as follows: starting from the interval [*ϕ*_0_, *ϕ*_1_], where *ϕ*_0_ = ^*m*^ *ϕ*_*low*_ and *ϕ*_0_ = ^*M*^ *ϕ*_low_, we calculated the value of *n*_*s*_ associated with the middle point of the interval (*ϕ*_1/2_). If *n*_*s*_ = 100, then *ϕ*_0_ was set to *ϕ*_1/2_. Otherwise, *ϕ*_1_ was set to *ϕ*_1/2_. This step was iterated until *ϕ*_1_ − *ϕ*_0_ < 10 ^− 3^.

A similar approach was used to estimate *ϕ*_*upp*_: the lower bound (^*m*^*ϕ*_*upp*_) was set as the value of *ϕ*_*D*_ among the previously explored points that produced the smallest non-zero *n*_*s*_. If one of those points produced an *n*_*s*_ equal to zero, then that point was set as the upper bound Otherwise, ^*M*^*ϕ*_*upp*_ = min x1.5 *ϕ*_ref_, 1000), whether the associated *n*_*s*_ is zero or not.

## Acknowledgements

The authors thank J. Cuesta and Iranzo’s group members for fruitful conversations, and L. Dudbridge for critical reading and editing of the manuscript. This work has been funded by the Agencia Estatal de Investigación of Spain (grant nos. PID2019-106618GA-I00 and CNS2023-145430 to JI), the Margarita Salas program of the Ministry of Universities of Spain (grant no. CT31/21 to JCE), the Ramón y Cajal program of the Spanish Ministry of Science (grant no. RYC-2017–22524 to JI), the Youth Employment Initiative of the European Social Fund through a junior postdoctoral contract from Comunidad de Madrid (grant no. PEJD-2019-POST/BIO-16377 to JCE), the Comunidad de Madrid through the call Research Grants for Young Investigators from Universidad Politécnica de Madrid (grant no. M190020074JIIS to JI), and the Severo Ochoa program for Centres of Excellence in R&D of the Agencia Estatal de Investigación of Spain (grant no. SEV-2016–0672 (2017–2021) to the CBGP).

## Notes

### Competing Interest Statement

The authors have declared no competing interest.

## References

1. Azam, F. and F. Malfatti, Microbial structuring of marine ecosystems. Nat Rev Microbiol, 2007. 5(10): p. 782–91.

2. Bardgett, R.D. and W.H. van der Putten, Belowground biodiversity and ecosystem functioning. Nature, 2014. 515(7528): p. 505–11.

3. McFall-Ngai, M., et al., Animals in a bacterial world, a new imperative for the life sciences. Proc Natl Acad Sci U S A, 2013. 110(9): p. 3229–36.

4. Falkowski, P.G., T. Fenchel, and E.F. Delong, The microbial engines that drive Earth’s biogeochemical cycles. Science, 2008. 320(5879): p. 1034–9.

5. Voolstra, C.R. and M. Ziegler, Adapting with Microbial Help: Microbiome Flexibility Facilitates Rapid Responses to Environmental Change. Bioessays, 2020. 42(7): p. e2000004.

6. Osburn, E.D., et al., Evaluating the role of bacterial diversity in supporting soil ecosystem functions under anthropogenic stress. ISME Commun, 2023. 3(1): p. 66.

7. Gilbert, J.A., et al., Current understanding of the human microbiome. Nat Med, 2018. 24(4): p. 392–400.

8. Smith, V.H. and D.W. Schindler, Eutrophication science: where do we go from here? Trends Ecol Evol, 2009. 24(4): p. 201–7.

9. Hillebrand, H. and U. Sommer, Diversity of benthic microalgae in response to colonization time and eutrophication. Aquatic Botany, 2000. 67(3): p. 221–236.

10. Lloyd-Price, J., et al., Multi-omics of the gut microbial ecosystem in inflammatory bowel diseases. Nature, 2019. 569(7758): p. 655–662.

11. Bai, X., et al., Engineering the gut microbiome. Nature Reviews Bioengineering, 2023. 1(9): p. 665–679.

12. Arif, I., M. Batool, and P.M. Schenk, Plant Microbiome Engineering: Expected Benefits for Improved Crop Growth and Resilience. Trends Biotechnol, 2020. 38(12): p. 1385–1396.

13. Rosado, P.M., et al., Marine probiotics: increasing coral resistance to bleaching through microbiome manipulation. ISME J, 2019. 13(4): p. 921–936.

14. Peixoto, R.S., et al., Harnessing the microbiome to prevent global biodiversity loss. Nat Microbiol, 2022. 7(11): p. 1726–1735.

15. Cavicchioli, R., et al., Scientists’ warning to humanity: microorganisms and climate change. Nat Rev Microbiol, 2019. 17(9): p. 569–586.

16. Albright, M.B.N., et al., Solutions in microbiome engineering: prioritizing barriers to organism establishment. ISME J, 2022. 16(2): p. 331–338.

17. May, R.M., Will a large complex system be stable? Nature, 1972. 238(5364): p. 413–4.

18. Landi, P., et al., Complexity and stability of ecological networks: a review of the theory. Population Ecology, 2018. 60(4): p. 319–345.

19. McCann, K.S., The diversity-stability debate. Nature, 2000. 405(6783): p. 228–33.

20. Larsen, P.E., S.M. Gibbons, and J.A. Gilbert, Modeling microbial community structure and functional diversity across time and space. FEMS Microbiol Lett, 2012. 332(2): p. 91–8.

21. Costello, E.K., et al., The application of ecological theory toward an understanding of the human microbiome. Science, 2012. 336(6086): p. 1255–62.

22. Coyte, K.Z., J. Schluter, and K.R. Foster, The ecology of the microbiome: Networks, competition, and stability. Science, 2015. 350(6261): p. 663–6.

23. Yonatan, Y., et al., Complexity-stability trade-off in empirical microbial ecosystems. Nat Ecol Evol, 2022. 6(6): p. 693–700.

24. Grilli, J., Macroecological laws describe variation and diversity in microbial communities. Nat Commun, 2020. 11(1): p. 4743.

25. Mataigne, V., et al., Microbial Systems Ecology to Understand Cross-Feeding in Microbiomes. Front Microbiol, 2021. 12: p. 780469.

26. Muller, E.E.L., et al., Using metabolic networks to resolve ecological properties of microbiomes. Current Opinion in Systems Biology, 2018. 8: p. 73–80.

27. Fisher, C.K. and P. Mehta, Identifying keystone species in the human gut microbiome from metagenomic timeseries using sparse linear regression. PLoS One, 2014. 9(7): p. e102451.

28. Stein, R.R., et al., Ecological modeling from time-series inference: insight into dynamics and stability of intestinal microbiota. PLoS Comput Biol, 2013. 9(12): p. e1003388.

29. Hatton, I.A., et al., Diversity begets stability: Sublinear growth and competitive coexistence across ecosystems. Science, 2024. 383(6688): p. eadg8488.

30. Allesina, S. and S. Tang, Stability criteria for complex ecosystems. Nature, 2012. 483(7388): p. 205–8.

31. Dougoud, M., et al., The feasibility of equilibria in large ecosystems: A primary but neglected concept in the complexity-stability debate. PLoS Comput Biol, 2018. 14(2): p. e1005988.

32. Stone, L., The feasibility and stability of large complex biological networks: a random matrix approach. Sci Rep, 2018. 8(1): p. 8246.

33. Stone, L., The stability of mutualism. Nat Commun, 2020. 11(1): p. 2648.

34. Haydon, D., Pivotal Assumptions Determining the Relationship between Stability and Complexity: An Analytical Synthesis of the Stability-Complexity Debate. The American Naturalist, 1994. 144(1): p. 14–29.

35. Qian, J.J. and E. Akcay, The balance of interaction types determines the assembly and stability of ecological communities. Nat Ecol Evol, 2020. 4(3): p. 356–365.

36. Arese Lucini, F., et al., Diversity increases the stability of ecosystems. PLoS One, 2020. 15(4): p. e0228692.

37. Gibbs, T., S.A. Levin, and J.M. Levine, Coexistence in diverse communities with higher-order interactions. Proc Natl Acad Sci U S A, 2022. 119(43): p. e2205063119.

38. Schuster, P. and K. Sigmund, Replicator dynamics. J Theor Biol, 1983. 100(3): p. 533–538.

39. Hofbauer, J. and K. Sigmund, Evolutionary games and population dynamics. 1998: Cambridge University Press.

40. Zelezniak, A., et al., Metabolic dependencies drive species co-occurrence in diverse microbial communities. Proc Natl Acad Sci U S A, 2015. 112(20): p. 6449–54.

41. Pacheco, A.R., M. Moel, and D. Segre, Costless metabolic secretions as drivers of interspecies interactions in microbial ecosystems. Nat Commun, 2019. 10(1): p. 103.

42. Morris, B.E., et al., Microbial syntrophy: interaction for the common good. FEMS Microbiol Rev, 2013. 37(3): p. 384–406.

43. Kelsic, E.D., et al., Counteraction of antibiotic production and degradation stabilizes microbial communities. Nature, 2015. 521(7553): p. 516–9.

44. Abrudan, M.I., et al., Socially mediated induction and suppression of antibiosis during bacterial coexistence. Proc Natl Acad Sci U S A, 2015. 112(35): p. 11054–9.

45. Levine, J.M., et al., Beyond pairwise mechanisms of species coexistence in complex communities. Nature, 2017. 546(7656): p. 56–64.

46. Mayfield, M.M. and D.B. Stouffer, Higher-order interactions capture unexplained complexity in diverse communities. Nat Ecol Evol, 2017. 1(3): p. 62.

47. Sanchez-Gorostiaga, A., et al., High-order interactions distort the functional landscape of microbial consortia. PLoS Biol, 2019. 17(12): p. e3000550.

48. Sundarraman, D., et al., Higher-Order Interactions Dampen Pairwise Competition in the Zebrafish Gut Microbiome. mBio, 2020. 11(5).

49. Chang, C.Y., et al., Emergent coexistence in multispecies microbial communities. Science, 2023. 381(6655): p. 343–348.

50. de Oliveira, V.M. and J.F. Fontanari, Random replicators with high-order interactions. Phys Rev Lett, 2000. 85(23): p. 4984–7.

51. Bairey, E., E.D. Kelsic, and R. Kishony, High-order species interactions shape ecosystem diversity. Nat Commun, 2016. 7: p. 12285.

52. Grilli, J., et al., Higher-order interactions stabilize dynamics in competitive network models. Nature, 2017. 548(7666): p. 210–213.

53. Wootton, J.T., The nature and consequences of indirect effects in ecological communities. Annual Review of Ecology, Evolution, and Systematics, 1994. 25(Volume 25): p. 443–466.

54. Shetty, S.A., et al., Dynamic metabolic interactions and trophic roles of human gut microbes identified using a minimal microbiome exhibiting ecological properties. ISME J, 2022. 16(9): p. 2144–2159.

55. Butler, S. and J.P. O’Dwyer, Stability criteria for complex microbial communities. Nat Commun, 2018. 9(1): p. 2970.

56. Sauve, A.M.C., C. Fontaine, and E. Thébault, Structure–stability relationships in networks combining mutualistic and antagonistic interactions. Oikos, 2014. 123(3): p. 378–384.

57. Foster, K.R. and T. Bell, Competition, not cooperation, dominates interactions among culturable microbial species. Curr Biol, 2012. 22(19): p. 1845–50.

58. Venturelli, O.S., et al., Deciphering microbial interactions in synthetic human gut microbiome communities. Mol Syst Biol, 2018. 14(6): p. e8157.

59. Goldford, J.E., et al., Emergent simplicity in microbial community assembly. Science, 2018. 361(6401): p. 469–474.

60. Coyte, K.Z. and S. Rakoff-Nahoum, Understanding Competition and Cooperation within the Mammalian Gut Microbiome. Curr Biol, 2019. 29(11): p. R538–R544.

61. Machado, D., et al., Polarization of microbial communities between competitive and cooperative metabolism. Nat Ecol Evol, 2021. 5(2): p. 195–203.

62. Ona, L., et al., Obligate cross-feeding expands the metabolic niche of bacteria. Nat Ecol Evol, 2021. 5(9): p. 1224–1232.

63. Duncker, K.E., Z.A. Holmes, and L. You, Engineered microbial consortia: strategies and applications. Microb Cell Fact, 2021. 20(1): p. 211.

64. Case, T.J., Invasion resistance arises in strongly interacting species-rich model competition communities. Proc Natl Acad Sci U S A, 1990. 87(24): p. 9610–4.

65. Roberts, A., The stability of a feasible random ecosystem. Nature, 1974. 251(5476): p. 607–608.

66. Donohue, I., et al., On the dimensionality of ecological stability. Ecol Lett, 2013. 16(4): p. 421–9.

67. Joseph, T.A., et al., Compositional Lotka-Volterra describes microbial dynamics in the simplex. PLoS Comput Biol, 2020. 16(5): p. e1007917.

68. Weiss, A.S., et al., In vitro interaction network of a synthetic gut bacterial community. ISME J, 2022. 16(4): p. 1095–1109.

69. Camacho-Mateu, J., et al., Sparse species interactions reproduce abundance correlation patterns in microbial communities. Proc Natl Acad Sci U S A, 2024. 121(5): p. e2309575121.

70. Freilich, S., et al., Competitive and cooperative metabolic interactions in bacterial communities. Nat Commun, 2011. 2: p. 589.

71. Clark, J.S., et al., Resolving the biodiversity paradox. Ecol Lett, 2007. 10(8): p. 647-59; discussion 659-62.

72. Holt, R.D., Emergent neutrality. Trends Ecol Evol, 2006. 21(10): p. 531–3.

73. Gianetto-Hill, C.M., et al., The Robogut: A Bioreactor Model of the Human Colon for Evaluation of Gut Microbial Community Ecology and Function. Curr Protoc, 2023. 3(4): p. e737.

74. Erez, A., et al., Nutrient levels and trade-offs control diversity in a serial dilution ecosystem. Elife, 2020. 9.

75. Gokhale, C.S. and A. Traulsen, Higher-order equivalence of Lotka-Volterra and replicator dynamics. bioRxiv, 2025: p. 2025.03.28.645916.

76. Hynes, W.F., et al., Bioprinting microbial communities to examine interspecies interactions in time and space. Biomedical Physics & Engineering Express, 2018. 4(5): p. 055010.

77. Blasche, S., et al., Metabolic cooperation and spatiotemporal niche partitioning in a kefir microbial community. Nat Microbiol, 2021. 6(2): p. 196–208.

