## Supplementary Information and Figures for "Stability and diversity of interactions in complex microbiomes"

### Stability and diversity of interactions in complex microbiomes
### Supplementary Information

Jorge Calle-Espinosa and Jaime Iranzo

**5 Supplementary Figures. 28**

**References 31**

### 1 Replicator equations for complex microbial communities.

For the sake of completeness, the specific formulation of the replicator equations analysed in the main body of the paper is stated again here. A system of $N$ species is considered. The frequency $x_i$ of each species in the population is given by the following replicator equations (equation 1.0.1):

$$\dot{x_i} = x_i\left(\Pi_i - \sum_{k=1}^{N}\Pi_k x_k\right) = x_i\left(\Pi_i - \overline{\Pi}\right) \tag{1.0.1}$$

where $\Pi_i$ represent the fitness of the species $i$ and $\overline{\Pi}$ the mean fitness of the population. The fitness term $\Pi_i$ (equation 1.0.2) depends on the interactions in which species $i$ is involved, including an inhibitory self-interaction term $-x_i$ that guarantees stability in the absence of other interactions:

$$\Pi_i = -x_i + \sum_{D=2}^{\infty}\sum^{\binom{N-1}{D-1}} {}_D\alpha_{i;k_1,k_2,...,k_{D-1}} x_{k_1}x_{k_2}...x_{k_{D-1}};\ k_j \neq i \tag{1.0.2}$$

where $D$ represents the order of the interaction ($i.e$ the number of species *simultaneously* involved in the interaction) and ${}_D\alpha_{i;k_1,k_2,...,k_{D-1}}$ represents the effect of species $k_1, k_2, ..., k_{D-1}$ over species $i$. Note that equation 1.0.2 assumes that no species affects the rest through terms with exponents different than one and that the focal species $i$ only affects its own fitness through the self inhibition term $-x_i$.

Each interaction can be characterized by the benefit or cost that it implies to each of the species involved. For example, pairwise interactions can be classified as cooperative (+/+), competitive (-/-), exploitative (+/-), commensalist (+/0) or ammensalist (0/-). In a similar fashion, higher order interactions can be classified according to the number of species receiving a benefit, a cost and/or nothing from the interplay. The model assesses all possible combinations of interaction orders $D$, proportions of types, and connectivities (understood as the fraction of all the possible order $D$ interactions in a given network, $C_D$) under the assumption of unstructured networks and independence of the interaction networks of different orders. To simplify the analysis, the magnitude of the interactions effects was restricted to a fixed value $\phi_D$ depending on the order of interaction (thus, ${}_D\alpha_{i;k_1,k_2,...,k_{D-1}}$ can only take values in the set $\{-\phi_D, 0, \phi_D\}$ depending on the case).

### 2 Sufficient conditions for the coexistence of all the species.

#### 2.1 The Kantorovich Theorem.

We used the Kantorovich Theorem to derive sufficient conditions for the existence of a stationary state comprising all species (see Argyros and Ren [2015] for a detailed presentation and proof of the theorem and Lecerf and Saadé [2015] for a survey of Kantorovich-like theorems). In particular, this proof uses the form of the Kantorovich Theorem presented in Corollary 5. Section 3.1 of Lecerf and Saadé [2015].

Kantorovich Theorem states sufficient conditions for the Newton–Raphson method to converge in a given convex set $D$. The Newton–Raphson method is a numerical method used to find roots of general vector valued function $F(x)$ defined in the Banach space $X$ from a starting guess $x_0$ using the information contained in the Jacobian $'F(x)$ of $F(x)$. The Kantorovich Theorem states that, given the following conditions:

$$0 \leq ||'F(x_0)^{-1}F(x_0)|| \leq \eta \tag{2.1.1}$$

$$||'F(x_0)^{-1}('F(x) -' F(y))|| \leq K||x-y|| \text{ for some } K>0 \text{ and for all } x,y \in D \tag{2.1.2}$$

$$\gamma = K\eta \leq 1/2 \tag{2.1.3}$$

$$B(x_0, r\eta) = \{x \in X, \, ||x-x_0|| < r\eta\} \subseteq D; \, r = \frac{2}{1+\sqrt{1-2\gamma}} \tag{2.1.4}$$

the Newton-Raphson method converges to a solution $x^* \in B(x_0, r\eta)$ of equation $F(x) = 0$.

To apply the Kantorovich Theorem, a proper function $F(x)$ needs to be built first. The objective here is to find stationary states that contain all the species in the system. According to equation 1.0.2:

$$\Pi_i - \overline{\Pi} = 0 \text{ for all } i \tag{2.1.5}$$

Equation 2.1.5 can be reformulated considering that in the stationary state $\overline{\Pi} = \frac{1}{N}\sum_{k=1}^{N}\Pi_k$:

$$\Pi_i - \frac{1}{N}\sum_{k=1}^{N}\Pi_k = 0 \text{ for all } i \tag{2.1.6}$$

Substituting the fitness values as defined by equation 1.0.2 and operating, we get to the following definition of $F(x)$:

$$F_i(x) = -x_i + \frac{1}{N}\sum_{k=1}^{N}x_k + \sum_{D=2}^{\infty}\left[\sum^{\binom{N-1}{D-1}} {}_D\alpha_{i;k_1,\ldots,k_{D-1}}x_{k_1}\ldots x_{k_{D-1}} - \frac{1}{N}\sum_{k=1}^{N}\sum^{\binom{N-1}{D-1}} {}_D\alpha_{k;k_1,\ldots,k_{D-1}}x_{k_1}\ldots x_{k_{D-1}}\right] \tag{2.1.7}$$

By definition, the set of $N$ equations defined in 2.1.7 is degenerate. To avoid that, and without loss of generality, the equation corresponding to the index $i = N$ is substituted by the following:

$$F_N(x) = \sum_{k=1}^{N}x_k - 1 \tag{2.1.8}$$

that represents the fact that the variables $x_i$ are fractions.

It is clear that a vector $x^*$ that fulfills $F_i(x^*) = 0$ is a stationary state of equation 1.0.1. To proof sufficient conditions for the existence of such stationary state, the Kantorovich Theorem will be applied using the system of equations defined in 2.1.7 and 2.1.8; the vector $x_0 = \frac{1}{N}\mathbb{1}$ (where $\mathbb{1}$ is the vector of ones) as the initial condition; and the infinity norm as the chosen norm. If with this selection of parameters conditions 2.1.1-2.1.4 are fulfilled and $B(x_0, r\eta) \subset B(x_0, \frac{1}{N})$ (that is, a solution exists that fulfills $0 < x_i < \frac{2}{N}$ for all species $i$), then it can be guaranteed that a stationary state $x^*$ with nonzero abundances of all species exists.

The Jacobian $'F(x)$ of $F(x)$ can be obtained analytically:

$$
\begin{gathered}
'F_{i,i}(x) = -1 + \frac{1}{N} - \sum_{D=2}^{\infty} \frac{1}{N} \sum_{k\neq i}^{N-1} \sum^{\binom{N-2}{D-2}} {}_{D}\alpha_{k;i,k_1,...,k_{D-2}} x_{k_1} ... x_{k_{D-2}} \\
'F_{i,j}(x) = \frac{1}{N} + \sum_{D=2}^{\infty} \left[ \sum^{\binom{N-2}{D-2}} {}_{D}\alpha_{i;j,k_1,...,k_{D-2}} x_{k_1} ... x_{k_{D-2}} - \frac{1}{N} \sum_{k\neq j}^{N-1} \sum^{\binom{N-2}{D-2}} {}_{D}\alpha_{k;j,k_1,...,k_{D-2}} x_{k_1} ... x_{k_{D-2}} \right] \\
'F_{N,j}(x) = 1
\end{gathered}
\tag{2.1.9}
$$

where the subindices $i, j$ take values in the set $\{1, 2, ..., N\}$.

Although the expressions of $F(x)$ and $'F(x)$ are a little cumbersome, when evaluated in $x_0 = \frac{1}{N}\mathbb{1}$ as required by the Kantorovich Theorem, they become much simpler. On one hand, $F(x_0)$ is:

$$
\begin{gathered}
F_i(x_0) = \sum_{D=2}^{\infty} \binom{N-1}{D-1} / N^{D-1} \left[ \langle {}_{D}\alpha_{i;\_} \rangle - \langle {}_{D}\alpha_{\_;\_} \rangle \right] \\
F_N(x_0) = 0
\end{gathered}
\tag{2.1.10}
$$

where $\langle {}_{D}\alpha_{i;\_} \rangle = \left( \sum^{\binom{N-1}{D-1}} {}_{D}\alpha_{i;k_1,...,k_{D-1}} \right) / \binom{N-1}{D-1}$ represents the mean effect of all sets of $D-1$ species on species $i$; and $\langle {}_{D}\alpha_{\_;\_} \rangle = \left( \sum_{k=1}^{N} \langle {}_{D}\alpha_{k;\_} \rangle \right) / N$ is the mean impact of all order $D$ interactions, excluding the self-inhibition term. The symbol $\langle \cdot \rangle$ denotes the mean; the $\alpha$ subindex **before** the semicolon indicates the **affected** species (all species if '_' is used); and the $\alpha$ subindices **after** the semicolon the set of species that **affect** the species indicated before the semicolon (again, '_' indicates all species sets). On the other hand, $'F(x_0)$ is:

$$
\begin{gathered}
'F_{i,i}(x_0) = -\left(1 - \frac{1}{N}\right) \left(1 + \sum_{D=2}^{\infty} \binom{N-2}{D-2} \langle {}_{D}\alpha_{\_;i,\_} \rangle / N^{D-2} \right) \\
'F_{i,j}(x_0) = \sum_{D=2}^{\infty} \binom{N-2}{D-2} / N^{D-2} \left[ \langle {}_{D}\alpha_{i;j,\_} \rangle - \langle {}_{D}\alpha_{\_;j,\_} \rangle \right] + \frac{1}{N} \left( 1 + \sum_{D=2}^{\infty} \binom{N-2}{D-2} / N^{D-2} \langle {}_{D}\alpha_{\_;j,\_} \rangle \right) \\
'F_{N,j}(x_0) = 1
\end{gathered}
\tag{2.1.11}
$$

where $\langle {}_{D}\alpha_{i;j,\_} \rangle = \left( \sum^{\binom{N-2}{D-2}} {}_{D}\alpha_{i;j,k_1,...,k_{D-2}} \right) / \binom{N-2}{D-2}$ indicates the mean effect on species $i$ of all sets of $D-1$ species containing species $j$; and $\langle {}_{D}\alpha_{\_;i,\_} \rangle = \left( \sum_{k\neq i}^{N-1} \langle {}_{D}\alpha_{k;i,\_} \rangle \right) / (N-1)$ represents the mean effect on all species of all sets of $D-1$ species containing species $i$.

#### 2.2 Dealing with $'F(x_0)$.

To verify conditions 2.1.1 and 2.1.2, the inverse of $'F(x_0)$ needs to be known. To deal with this calculation, let us assume that matrix $'F(x_0)$ is decomposed in matrices $L$ (assumed invertible) and $\Delta$.

Under this assumption, $'F(x_0)^{-1}$ can be reformulated as follows:

$$'F(x_0)^{-1} = [L + \Delta]^{-1} = [I + L^{-1}\Delta]^{-1}L^{-1} \tag{2.2.1}$$

A proper decomposition of $'F(x_0)^{-1}$ is given by the following matrices:

$$L = \begin{bmatrix} -(1-\frac{1}{N})\beta & \beta/N & \cdots & \beta/N & \beta/N \\ \beta/N & -(1-\frac{1}{N})\beta & \cdots & \beta/N & \beta/N \\ \vdots & \vdots & \ddots & \vdots & \vdots \\ \beta/N & \beta/N & \cdots & -(1-\frac{1}{N})\beta & \beta/N \\ 1 & 1 & \cdots & 1 & 1 \end{bmatrix} \tag{2.2.2}$$

$$\text{with } \beta = 1 + \sum_{D=2}^{\infty} \binom{N-2}{D-2} / N^{D-2} \mathbb{E}\big[\langle_D \alpha_{\text{-};\text{-}} \rangle\big]$$

$$\Delta = \begin{bmatrix} \Delta_{1,1} & \Delta_{1,2} & \cdots & \Delta_{1,N-1} & \Delta_{1,N} \\ \Delta_{2,1} & \Delta_{2,2} & \cdots & \Delta_{2,N-1} & \Delta_{2,N} \\ \vdots & \vdots & \ddots & \vdots & \vdots \\ \Delta_{N-1,1} & \Delta_{N-1,2} & \cdots & \Delta_{N-1,N-1} & \Delta_{N-1,N} \\ 0 & 0 & \cdots & 0 & 0 \end{bmatrix}$$

$$\text{with } \Delta_{i,i} = \sum_{D=2}^{\infty} -\left(1 - \frac{1}{N}\right)\binom{N-2}{D-2} / N^{D-2} \Big[\langle_D \alpha_{\text{-};i,\text{-}} \rangle - \mathbb{E}\big[\langle_D \alpha_{\text{-};\text{-}} \rangle\big]\Big]$$

$$\text{and } \Delta_{i,j} = \sum_{D=2}^{\infty} \binom{N-2}{D-2} / N^{D-2} \Big[\langle_D \alpha_{i;j,\text{-}} \rangle - \langle_D \alpha_{\text{-};j,\text{-}} \rangle\Big] + \frac{1}{N} \sum_{D=2}^{\infty} \binom{N-2}{D-2} / N^{D-2} \Big[\langle_D \alpha_{\text{-};j,\text{-}} \rangle - \mathbb{E}\big[\langle_D \alpha_{\text{-};\text{-}} \rangle\big]\Big] \tag{2.2.3}$$

There are several reasons that justify the above decomposition. First, the elements of matrix $\Delta$ are obtained from deviations of the effect on the community of a particular species with respect to the expected effect among species ($\langle_D \alpha_{\text{-};i,\text{-}} \rangle - \mathbb{E}\big[\langle_D \alpha_{\text{-};\text{-}} \rangle\big]$) and/or the deviations of the effect of a particular species $j$ on species $i$ with respect to the mean of the effect of species $j$ over the rest ($\langle_D \alpha_{i;j,\text{-}} \rangle - \langle_D \alpha_{\text{-};j,\text{-}} \rangle$). Therefore, if every species affects all the other species in the community in the same way, the matrix $\Delta$ becomes a matrix of zeros. Under this perspective, $L$ represents a limit matrix towards which $'F(x_0)$ tends when all the species in the system are equivalent in terms of interactions.

From a more technical perspective, the limit matrix $L$ is very easy to invert:

$$L^{-1} = \begin{bmatrix} -1/\beta & 0 & \cdots & 0 & 1/N \\ 0 & -1/\beta & \cdots & 0 & 1/N \\ \vdots & \vdots & \ddots & \vdots & \vdots \\ 0 & 0 & \cdots & -1/\beta & 1/N \\ 1/\beta & 1/\beta & \cdots & 1/\beta & 1/N \end{bmatrix} \tag{2.2.4}$$

$$\text{with } \beta = 1 + \sum_{D=2}^{\infty} \binom{N-2}{D-2} / N^{D-2} \mathbb{E}\left[\langle_D \alpha_{\text{-};\text{-}} \rangle\right]$$

and the form of $L^{-1}$ makes the products $L^{-1}M$ very simple in general, but particularly for the matrices and vectors that are relevant for the application of the Kantorovich Theorem, specifically the vector $F(x_0)$ (condition 2.1.1) and the matrices $'F(x) -' F(y)$ (condition 2.1.2) and $\Delta$ (used to estimate an upper bound for $\left|L^{-1}F(x_0)\right|_\infty$ in particular scenarios). The corresponding products are:

$$L^{-1}\Delta = -1/\beta \begin{bmatrix} \Delta_{1,1} & \Delta_{1,2} & \cdots & \Delta_{1,N-1} & \Delta_{1,N} \\ \Delta_{2,1} & \Delta_{2,2} & \cdots & \Delta_{2,N-1} & \Delta_{2,N} \\ \vdots & \vdots & \ddots & \vdots & \vdots \\ \Delta_{N-1,1} & \Delta_{N-1,2} & \cdots & \Delta_{N-1,N-1} & \Delta_{N-1,N} \\ \Delta_{N,1} & \Delta_{N,2} & \cdots & \Delta_{N,N-1} & \Delta_{N,N} \end{bmatrix}$$

$$\text{with } \beta = 1 + \sum_{D=2}^{\infty} \binom{N-2}{D-2} / N^{D-2} \mathbb{E}\left[\langle_D \alpha_{\text{-};\text{-}} \rangle\right]$$

$$\text{and } \Delta_{i,i} = \sum_{D=2}^{\infty} -\left(1 - \frac{1}{N}\right) \binom{N-2}{D-2} / N^{D-2} \left[\langle_D \alpha_{\text{-};i,\text{-}} \rangle - \mathbb{E}\left[\langle_D \alpha_{\text{-};\text{-}} \rangle\right]\right]$$

$$\text{and } \Delta_{i,j} = \sum_{D=2}^{\infty} \binom{N-2}{D-2} / N^{D-2} \left[\langle_D \alpha_{i;j,\text{-}} \rangle - \langle_D \alpha_{\text{-};j,\text{-}} \rangle\right] + \frac{1}{N} \sum_{D=2}^{\infty} \binom{N-2}{D-2} / N^{D-2} \left[\langle_D \alpha_{\text{-};j,\text{-}} \rangle - \mathbb{E}\left[\langle_D \alpha_{\text{-};\text{-}} \rangle\right]\right] \tag{2.2.5}$$

$$L^{-1}F(x_0)_i = -\sum_{D=2}^{\infty} \binom{N-1}{D-1} / N^{D-1} \left[\langle_D \alpha_{i;\text{-}} \rangle - \langle_D \alpha_{\text{-};\text{-}} \rangle\right] / \beta \quad \text{for all } i \in \{1, 2, \cdots, N\} \tag{2.2.6}$$

$$\text{with } \beta = 1 + \sum_{D=2}^{\infty} \binom{N-2}{D-2} / N^{D-2} \mathbb{E}\left[\langle_D \alpha_{\text{-};\text{-}} \rangle\right]$$

$$L^{-1}[{}'F(x) -{}' F(y)] = -1/\beta \begin{bmatrix} \Delta' F_{1,1} & \Delta' F_{1,2} & \cdots & \Delta' F_{1,N-1} & \Delta' F_{1,N} \\ \Delta' F_{2,1} & \Delta' F_{2,2} & \cdots & \Delta' F_{2,N-1} & \Delta' F_{2,N} \\ \vdots & \vdots & \ddots & \vdots & \vdots \\ \Delta' F_{N-1,1} & \Delta' F_{N-1,2} & \cdots & \Delta' F_{N-1,N-1} & \Delta' F_{N-1,N} \\ \Delta' F_{N,1} & \Delta' F_{N,2} & \cdots & \Delta' F_{N,N-1} & \Delta' F_{N,N} \end{bmatrix}$$

$$\text{with } \beta = 1 + \sum_{D=2}^{\infty} \binom{N-2}{D-2} / N^{D-2} \mathbb{E}\left[\langle {}_D\alpha_{-;-} \rangle\right]$$

$$\text{and } \Delta' F_{i,i} = \sum_{D=2}^{\infty} \frac{1}{N} \sum_{k \neq i}^{N-1} \sum_{k_l \neq i,k}^{\binom{N-2}{D-2}} {}_D\alpha_{k;i,k_1,\ldots,k_{D-2}} (x_{k_1} \ldots x_{k_{D-2}} - y_{k_1} \ldots y_{k_{D-2}}) \tag{2.2.7}$$

$$\text{and } \Delta' F_{i,j} = \sum_{D=2}^{\infty} \Bigg[ \sum_{k_l \neq i,j}^{\binom{N-2}{D-2}} {}_D\alpha_{i;j,k_1,\ldots,k_{D-2}} (x_{k_1} \ldots x_{k_{D-2}} - y_{k_1} \ldots y_{k_{D-2}}) -$$
$$- \frac{1}{N} \sum_{k \neq j}^{N-1} \sum_{k_l \neq j,k}^{\binom{N-2}{D-2}} \langle {}_D\alpha_{k;j,k_1,\ldots,k_{D-2}} \rangle (x_{k_1} \ldots x_{k_{D-2}} - y_{k_1} \ldots y_{k_{D-2}}) \Bigg]$$

In all cases, the product $L^{-1}M$ is just $M$ multiplied by a constant $-1/\beta$, but with the $N^{th}$ row changed from a deviant one with respect to the rest of the rows, to a row which is consistent with the rest (see, for example, that the row of zeros in matrix $\Delta$ is substituted by a row of elements $\Delta_{N,k}$, which has a form analogous to the rest of the rows). Given these products, the relevant norms for the assessment of the Kantorovich Theorem can be estimated.

#### 2.3 Calculation of $\eta$.

The first constant to be calculated for the application of the Kantorovich Theorem is $\eta$ (see conditions 2.1.1 and 2.1.3 and equation 2.1.4). To do so, let us start with the following inequality:

$$\left| {}'F(x_0)^{-1} F(x_0) \right|_\infty = \left| [I + L^{-1}\Delta]^{-1} L^{-1} F(x_0) \right|_\infty \leq \left| [I + L^{-1}\Delta]^{-1} \right|_\infty \left| L^{-1} F(x_0) \right|_\infty \tag{2.3.1}$$

By replacing the value of $\left| L^{-1} F(x_0) \right|_\infty$ with the upper bound derived in subsection 4.3:

$$\left| {}'F(x_0)^{-1} F(x_0) \right|_\infty \leq \left| [I + L^{-1}\Delta]^{-1} \right| \frac{\sqrt{2log(N)}}{|\beta|} \left( \sum_{D=2}^{\infty} \binom{N-1}{D-1} / N^{2(D-1)} \phi_D^2 C_D \left[ 1 - C_D \big(2P_D - 1\big)^2 \right] \right)^{1/2}$$
$$\text{with } \beta = 1 + \sum_{D=2}^{\infty} \phi_D C_D \big(2P_D - 1\big) \binom{N-2}{D-2} / N^{D-2} \tag{2.3.2}$$

Having an upper bound for $\left| L^{-1} F(x_0) \right|_\infty$, only an upper bound for $\left| [I + L^{-1}\Delta]^{-1} \right|_\infty$ needs to be calculated for defining $\eta$. However, an explicit tight upper bound of $\left| [I + L^{-1}\Delta]^{-1} \right|_\infty$ is difficult to obtain in general. A typical case that allows for an easy calculation of an upper bound of the norm of the inverse of a of matrix occurs when the matrix is strictly diagonally dominant (that is, the absolute value of the $i^{th}$ diagonal element is greater than the sum of the absolute values of the rest of the $i^{th}$ row, $|a_{i,i}| > \sum_{j\neq i} |a_{i,j}|$). Assuming this is the case for $\Big|[I + L^{-1}\Delta]^{-1}\Big|_\infty$, the Varah bound (Varah [1975]) can be used (see subsection 4.4), yielding a closed form of this upper bound (see inequalities 4.4.2 and 4.3.2). Unfortunately, this upper bound seems to be useful only for interaction networks dominated by order $\geq 5$ interaction, especially for large $N$. In these particular cases, $\Big|[I + L^{-1}\Delta]^{-1}\Big|_\infty \to 1$ as $N \to \infty$. In the rest of scenarios, the value of $\Big|[I + L^{-1}\Delta]^{-1}\Big|_\infty$ needs to be calculated numerically. In consequence, from now on, the term $\Big|[I + L^{-1}\Delta]^{-1}\Big|_\infty$ will be kept in the corresponding equations. In line with that, an upper bound for $\eta$ can be defined as:

$$\eta \leq \Big|[I + L^{-1}\Delta]^{-1}\Big|_\infty \frac{\sqrt{2log(N)}}{|\beta|}\left(\sum_{D=2}^{\infty}\binom{N-1}{D-1}/N^{2(D-1)}\phi_D^2 C_D\left[1 - C_D\big(2P_D - 1\big)^2\right]\right)^{1/2}$$
$$\text{with } \beta = 1 + \sum_{D=2}^{\infty}\phi_D C_D\big(2P_D - 1\big)\binom{N-2}{D-2}/N^{D-2} \tag{2.3.3}$$

which in the limit of $N \to \infty$ becomes:

$$\lim_{N\to\infty}\eta \leq \Big|[I + L^{-1}\Delta]^{-1}\Big|_\infty \frac{\sqrt{2log(N)}}{|\beta|}\left(\sum_{D=2}^{\infty}N^{1-D}\frac{\phi_D^2 C_D}{(D-1)!}\left[1 - C_D\big(2P_D - 1\big)^2\right]\right)^{1/2}$$
$$\text{with } \beta = 1 + \sum_{D=2}^{\infty}\frac{\phi_D C_D\big(2P_D - 1\big)}{(D-2)!} \tag{2.3.4}$$

#### 2.4 Calculation of $K$.

Having the value of $\eta$, only the constant $K$ remains to be estimated. To do so, the expression for the inverse matrix ${}'F(x_0)^{-1} \sim [I + L^{-1}\Delta]^{-1}L^{-1}$ presented in 2.2.1 will be used again. Under this assumption, an upper bound of $K$ is:

$$K|x - y|_\infty \leq \Big|[I + L^{-1}\Delta]^{-1}\Big|_\infty \Big|L^{-1}\big({}'F(x) -{}' F(y)\big)\Big|_\infty \tag{2.4.1}$$

The value of the norm $\Big|[I + L^{-1}\Delta]^{-1}\Big|_\infty$ is discussed during the derivation of $\eta$ and in subsection 4.4. Unfortunately, the strategies used until now no longer work to estimate the second term. However, a probably coarse upper bound can be obtained by bounding the absolute values of the elements of the matrix $L^{-1}\big({}'F(x) -{}' F(y)\big)$ (see equation 2.2.7) as follows:

$$|\Delta' F_{i,i}| = \frac{1}{|\beta|}\left|\sum_{D=2}^{\infty}\frac{1}{N}\sum_{k\neq i}^{N-1}\sum_{k_l\neq i,k}^{\binom{N-2}{D-2}} {}_D\alpha_{k;i,k_1,\dots,k_{D-2}}(x_{k_1}...x_{k_{D-2}} - y_{k_1}...y_{k_{D-2}})\right| \leq$$

$$\leq \sum_{D=2}^{\infty}\frac{\phi_D}{|\beta|}\left(1-\frac{D-1}{N}\right)\Big|\sum_{k_l\neq i}^{\binom{N-1}{D-2}} x_{k_1}...x_{k_{D-2}} - y_{k_1}...y_{k_{D-2}}\Big|$$

$$\begin{aligned}|\Delta' F_{i,j}| = \frac{1}{|\beta|}\Bigg|\sum_{D=2}^{\infty}\Bigg[\sum_{k_l\neq i,j}^{\binom{N-2}{D-2}} {}_D\alpha_{i;j,k_1,\dots,k_{D-2}}(x_{k_1}...x_{k_{D-2}} - y_{k_1}...y_{k_{D-2}})-\\ -\frac{1}{N}\sum_{k\neq j}^{N-1}\sum_{k_l\neq j,k}^{\binom{N-2}{D-2}}\langle {}_D\alpha_{k;j,k_1,\dots,k_{D-2}}\rangle(x_{k_1}...x_{k_{D-2}} - y_{k_1}...y_{k_{D-2}})\Bigg]\Bigg| \leq\end{aligned} \quad (2.4.2)$$

$$\sum_{D=2}^{\infty}\frac{\phi_D}{|\beta|}\left[\Big|\sum_{k_l\neq i,j}^{\binom{N-2}{D-2}} |x_{k_1}...x_{k_{D-2}} - y_{k_1}...y_{k_{D-2}}\Big| + \left(1-\frac{D-1}{N}\right)\Big|\sum_{k_l\neq j}^{\binom{N-1}{D-2}} x_{k_1}...x_{k_{D-2}} - y_{k_1}...y_{k_{D-2}}\Big|\right]$$

$$\text{with } \beta = 1 + \sum_{D=2}^{\infty}\phi_D C_D(2P_D-1)\binom{N-2}{D-2}/N^{D-2}$$

Giving these bounds, the term $\frac{\left|L^{-1}({}'F(x) -' F(y))\right|_\infty}{|x-y|_\infty} \leq K$ can be bounded as follows:

$$\begin{aligned}\frac{\left|L^{-1}({}'F(x) -' F(y))\right|_\infty}{|x-y|_\infty} &\leq \max_{i,x,y}\Bigg[\sum_{D=2}^{\infty}\frac{\phi_D}{|\beta|}\frac{\left|\sum_{j\neq i}^{N-1}\sum_{k_l\neq i,j}^{\binom{N-2}{D-2}} x_{k_1}...x_{k_{D-2}} - y_{k_1}...y_{k_{D-2}}\right|}{|x-y|_\infty} +\\ &+\frac{\phi_D}{|\beta|}\left(1-\frac{D-1}{N}\right)\frac{\left|\sum_{j=1}^{N}\sum_{k_l\neq j}^{\binom{N-1}{D-2}} x_{k_1}...x_{k_{D-2}} - y_{k_1}...y_{k_{D-2}}\right|}{|x-y|_\infty}\Bigg] =\\ &= \max_{i,x,y}\Bigg[\sum_{D=2}^{\infty}\frac{\phi_D}{|\beta|}(N-D+1)\frac{\left|\sum_{k_l\neq i}^{\binom{N-1}{D-2}} x_{k_1}...x_{k_{D-2}} - y_{k_1}...y_{k_{D-2}}\right|}{|x-y|_\infty} +\\ &+\frac{\phi_D}{|\beta|}\left(1-\frac{D-1}{N}\right)(N-D+2)\Big|\frac{\sum^{\binom{N}{D-2}} x_{k_1}...x_{k_{D-2}} - y_{k_1}...y_{k_{D-2}}}{|x-y|_\infty}\Big|\Bigg] \leq\\ &\leq \sum_{D=2}^{\infty}\frac{\phi_D}{|\beta|}(N-D+1)\max_{i,x,y}\left[\frac{\left|\sum_{k_l\neq i}^{\binom{N-1}{D-2}} x_{k_1}...x_{k_{D-2}} - y_{k_1}...y_{k_{D-2}}\right|}{|x-y|_\infty}\right] +\\ &+\frac{\phi_D}{|\beta|}\left(1-\frac{D-1}{N}\right)(N-D+2)\max_{x,y}\left[\frac{\left|\sum^{\binom{N}{D-2}} x_{k_1}...x_{k_{D-2}} - y_{k_1}...y_{k_{D-2}}\right|}{|x-y|_\infty}\right]\\ &\text{with } \beta = 1 + \sum_{D=2}^{\infty}\phi_D C_D(2P_D-1)\binom{N-2}{D-2}/N^{D-2}\end{aligned} \quad (2.4.3)$$

In subsection 4.6 the upper bounds $\max_{x,y}\left[\frac{\left|\sum^{\binom{N}{D-2}} x_{k_1}...x_{k_{D-2}} - y_{k_1}...y_{k_{D-2}}\right|}{|x-y|_\infty}\right] \leq 1/(D-3)!$ (zero if

$D \leq 3$) and $\max_{i,x,y}\left[\frac{\left|\sum_{k_l\neq i}^{\binom{N-1}{D-2}} x_{k_1}\ldots x_{k_{D-2}} - y_{k_1}\ldots y_{k_{D-2}}\right|}{|x-y|_\infty}\right] \leq 2/(D-3)!$ (zero if $D=2$ and one if $D=3$) were proven, allowing, along with inequalities 2.4.1 and 2.4.3, to formulate a closed expression for $K$:

$$K \leq \left|[I+L^{-1}\Delta]^{-1}\right|_\infty \left[\frac{\phi_3}{|\beta|}(N-2) + \sum_{D=4}^{\infty}\frac{\phi_D}{|\beta|(D-3)!}(N-D+1)\left(3-\frac{D-2}{N}\right)\right]$$
$$\text{with } \beta = 1 + \sum_{D=2}^{\infty}\phi_D C_D\left(2P_D-1\right)\binom{N-2}{D-2}/N^{D-2} \tag{2.4.4}$$

which in the limit of $N \to \infty$ becomes:

$$\lim_{N\to\infty} K \leq \left|[I+L^{-1}\Delta]^{-1}\right|_\infty \left(\frac{\phi_3}{|\beta|} + \sum_{D=4}^{\infty}\frac{3\phi_D}{|\beta|(D-3)!}\right)N$$
$$\text{with } \beta = 1 + \sum_{D=2}^{\infty}\frac{\phi_D C_D\left(2P_D-1\right)}{(D-2)!} \tag{2.4.5}$$

#### 2.5 Application of the Kantorovich Theorem.

With the boundaries of $\eta$ (inequality 2.3.3) and $K$ (inequality 2.4.4) defined, now the main condition (inequality 2.1.3) for the application of the Kantorovich Theorem can be assessed. The first important conclusion that arises from these boundaries is that, for high values of $N$, the boundaries are dominated by the smallest order $D_{min}$ with $\phi_{D_{min}} \neq 0$ unless the strength of interaction $\phi_D$ of associated to other orders are conveniently scaled with $N$. This is true even when the upper bound of $\left|[I+L^{-1}\Delta]^{-1}\right|_\infty$ cannot be estimated due to the behaviour of its lower bound with respect to $N$ and $D$ (see subsection 4.4).

If only one order is considered, the expressions of $\eta$ and $K$ are significantly simpler. In this cases, for minimum order $D > 2$ and assuming $N \to \infty$, the condition $\gamma = \eta K \leq 1/2$ takes the following form:

$$\gamma \leq 2\kappa\left|[I+L^{-1}\Delta]^{-1}\right|_\infty^2 \frac{\sqrt{2log(N)}}{\beta^2(D-3)!}\frac{\phi_D^2 C_D^{1/2}}{(D-1)!^{1/2}}\left[1-C_D\left(2P_D-1\right)^2\right]^{1/2} N^{(3-D)/2}$$
$$\text{with } \beta = 1 + \sum_{D=2}^{\infty}\frac{\phi_D C_D\left(2P_D-1\right)}{(D-2)!}, \ \kappa = \begin{cases}1 \text{ if } D=3\\ 3 \text{ if } D\geq 4\end{cases}, \text{ and } D>2 \tag{2.5.1}$$

According to inequality 2.5.1 and taking into account that $\left|[I+L^{-1}\Delta]^{-1}\right|_\infty \geq 1$, for order 3, $\gamma$ increases with $N$ at least as $\sqrt{2log(N)}$. Moreover, when the upper bound of $[I+L^{-1}\Delta]^{-1}\Big|_\infty$ (see subsection 4.4) can be applied, the value of $\gamma$ increases with $N$ as $\sqrt{2log(N)}N$. Thus, the possibility of application of the Kantorovich Theorem in this cases is limited to particular networks (small $\phi_3$ and/or extreme $C_3$ and $P_3$). However, for $D > 4$, $\gamma$ decreases with $N$ the faster the higher the order considered. Thus, in these scenarios, the Kantorovich Theorem can be usually applied, except in systems with small $N$, high $\phi_D$ (only in some situations) and/or values of $\phi_D C_D\left(2P_D-1\right)\binom{N-2}{D-2}/N^{D-2}$ or $\phi_D C_D\left(2P_D-1\right)\binom{N-1}{D-1}/N^{D-1}$ close to one. The particular case of $D=4$ requires a detailed discussion. First of all, if the $[I+L^{-1}\Delta]^{-1}\Big|_\infty$ increases with less than $N^{1/4}$, then the conclusions for orders greater than 4 can be applied: increasing $N$ allows the fulfillment of the conditions of the Kantorovich Theorem. This is compatible with the lower bound of $[I + L^{-1}\Delta]^{-1}\Big|_{\infty}$ tending to 1 as $N$ tends to $\infty$ (see subsection 4.4), but the lack of a proper upper bound when $I + L^{-1}\Delta$ is not strictly diagonally dominant impedes making any solid statement regarding how $[I+L^{-1}\Delta]^{-1}\Big|_{\infty}$ scales with $N$.

For order 2 networks alone, the function $F(x)$ is linear and thus the Kantorovich Theorem automatically applies because $K$ takes a value of 0. Unfortunately, when mixtures of orders are considered, the presence of order 2 interactions induces an increase of $\gamma$ proportional to more than $N^{1/2}$. In these situations, as in the case of order 3, only interaction networks with small $\phi_2$ and/or extreme $C_2$ and $P_2$ are susceptible to the application of the Kantorovich Theorem.

The problems with the application of the Kantorovich Theorem to order 2 and order 3 interaction networks can be compensated by the higher order interactions if their intensity $\phi_D$ is proportional to some positive power of $N$ thanks to the presence of the term $\phi_D C_D \big(2P_D - 1\big)\binom{N-2}{D-2}/N^{D-2}$. However, it is important to note that increasing $\phi_D$ with some power of $N$ also affects $\gamma$ in other ways, specially reducing the impact of the self-inhibition in the system.

Let us now focus on the cases in which the Kantorovich Theorem can be applied. As described previously, if the conditions of the theorem are fulfilled, then a solution exists in the ball $B(x_0, r\eta) = \{x \in X, |x - x_0|_\infty < r\eta\}$, where $x_0$ is the vector with all components equal to $1/N$, that is, the homogeneous community. Considering that, for a stationary state with all species present to exist:

$$r\eta \leq 1/N \tag{2.5.2}$$

As shown in condition 2.1.4, $r = \frac{2}{1+\sqrt{1-2\gamma}}$, so if $\gamma \leq 1/2$ as required by the theorem, $r$ takes values between 1 and 2. For simplicity, let us assume that $r = 2$, so the complex expression of $\gamma$ does not need to be used. Then, by inequality 2.3.3, the sufficient condition for existence is:

$$\begin{gathered}
2N\Big|[I + L^{-1}\Delta]^{-1}\Big|_{\infty} \frac{\sqrt{2log(N)}}{|\beta|}\left(\sum_{D=2}^{\infty}\binom{N-1}{D-1}/N^{2(D-1)}\phi_D^2 C_D\Big[1 - C_D\big(2P_D - 1\big)^2\Big]\right)^{1/2} \leq 1 \\
\text{with } \beta = 1 + \sum_{D=2}^{\infty}\phi_D C_D\big(2P_D - 1\big)\binom{N-2}{D-2}/N^{D-2}
\end{gathered} \tag{2.5.3}$$

that for large $N$ becomes:

$$\begin{gathered}
2\Big|[I + L^{-1}\Delta]^{-1}\Big|_{\infty} \frac{\sqrt{2log(N)}}{|\beta|}\left(\sum_{D=2}^{\infty} N^{3-D}\frac{\phi_D^2 C_D}{(D-1)!}\Big[1 - C_D\big(2P_D - 1\big)^2\Big]\right)^{1/2} \leq 1 \\
\text{with } \beta = 1 + \sum_{D=2}^{\infty}\frac{\phi_D C_D\big(2P_D - 1\big)}{(D-2)!}
\end{gathered} \tag{2.5.4}$$

Condition 2.5.3 is identical to the one required for the application of the Kantorovich Theorem (in the limit $N \to \infty$) but for two factors, being one $\Big|[I + L^{-1}\Delta]^{-1}\Big|_{\infty}$ and the other independent of $N$. Thus, almost all the considerations regarding the application of the Kantorovich Theorem can be immediately extrapolated to condition 2.5.3. However, it is important to emphasize one difference: in the case of networks with significant presence of order 2 and 3 interactions, the destabilizing effect of $N$ is expected to be lower due to $\left|[I + L^{-1}\Delta]^{-1}\right|_\infty$ not being squared in condition 2.5.3, making the compensation of that effect *via* higher orders easier.

In summary, we conclude that in networks characterized by interactions of order greater than 4 (and probably 4), increasing the number of species $N$ facilitates the existence of a stationary state with nonzero abundances of all the $N$ species. Not only that, but the greater the $N$ the closer this stationary state is to a homogeneous composition with all species present in the same proportion. This is due to the fact that the smaller the left side of inequality 2.5.3, the smaller is the radius $r\eta$ of the ball $B(x_0, r\eta) = \{x \in X, |x - x_0|_\infty < r\eta\}$ and, thus, the closer the solution to the chosen $x_0$, which is the homogeneous community in this case.

If lower interaction orders are dominant, these sufficient conditions are of limited use. The reason is that, in those cases, the range of interaction networks that fulfill the different conditions shrinks as $N$ increases. This happens even if only the less strict condition 2.5.3 is considered. Although the Kantorovich Theorem only states sufficient conditions for the existence of solutions, the destabilizing effect of $N$ in low-order interaction networks has been confirmed with simulations, as described in the main text. Thus, the impossibility of applying the Kantorovich Theorem and/or the absence of a guaranteed solution with all species present in these cases responds, at least partially, to the fact that these solutions do not exist in reality.

Finally, and whichever order considered, the networks that are most prone to fulfill the Kantorovich Theorem and produce stationary states with nonzero fractions of all the species are the following: (i) almost unconnected networks, and (ii) densely connected and highly polarized networks (almost pure cooperative or competitive cliques).

### 3 Conditions for stability.

#### 3.1 Analytical derivation

In this section, we derive conditions for the local stability of stationary states under particular assumptions. For a stationary state to be stable, the eigenvalues of the Jacobian of the system of differential equations, evaluated at that stationary state, must be negative. In the present case, the Jacobian of system 1.0.1 is:

$$\begin{aligned} J_{i,i} &= \left(\Pi_i - \overline{\Pi}\right) + x_i^{SS}\left(\frac{\partial \Pi_i}{\partial x_i} - \frac{\partial \overline{\Pi}}{\partial x_i}\right) \\ J_{i,j} &= x_i^{SS}\left(\frac{\partial \Pi_i}{\partial x_j} - \frac{\partial \overline{\Pi}}{\partial x_j}\right) \end{aligned} \tag{3.1.1}$$

where $x_i^{SS}$ represents the frequency of species $i$ at the stationary state. If a stationary state including all species exists, then all fitness values are the same, that is $\Pi_i = \overline{\Pi}$ for all $i$. Thus, the Jacobian for the stationary states becomes:

$$\begin{aligned} J_{i,i} &= x_i^{SS}\left(\frac{\partial \Pi_i}{\partial x_i} - \frac{\partial \overline{\Pi}}{\partial x_i}\right) \\ J_{i,j} &= x_i^{SS}\left(\frac{\partial \Pi_i}{\partial x_j} - \frac{\partial \overline{\Pi}}{\partial x_j}\right) \end{aligned} \tag{3.1.2}$$

By substituting the partial derivatives from equation 1.0.2, the following expressions are obtained:

$$J_{i,i} = x_i^{SS}\Bigg[x_i^{SS} - \overline{\Pi} - 1 - \sum_{k\neq i}^{N-1} x_k^{SS} \sum_{D=2}^{\infty} \sum^{\binom{N-2}{D-2}} {}_D\alpha_{k;i,k_2,\ldots,k_{D-2}} x_{k_1}^{SS} x_{k_2}^{SS} \ldots x_{k_{D-2}}^{SS}\Bigg]$$

$$J_{i,j} = x_i^{SS}\Bigg[x_j^{SS} - \overline{\Pi} + \sum_{D=2}^{\infty} \sum^{\binom{N-2}{D-2}} \Big({}_D\alpha_{i;j,k_2,\ldots,k_{D-2}} x_{k_1}^{SS} x_{k_2}^{SS} \ldots x_{k_{D-2}}^{SS} - \sum_{k\neq j}^{N-1} x_k^{SS}\, {}_D\alpha_{k;j,k_2,\ldots,k_{D-2}} x_{k_1}^{SS} x_{k_2}^{SS} \ldots x_{k_{D-2}}^{SS}\Big)\Bigg] \quad (3.1.3)$$

Because $J^T\mathbb{1} = -\overline{\Pi}\mathbb{1}$ (where the $T$ indicates the transpose and $\mathbb{1}$ is a column vector of ones), it is clear that $-\overline{\Pi}$ is an eigenvalue of 3.1.3 with $u = \mathbb{1}$ as the associated eigenvector. Note that this eigenvalue just indicates that if the mean fitness at the stationary state is not positive, the whole community will become extinct. However, if all species are allowed to grow independently at a rate $r$, the replicator equations will remain identical to those in 1.0.1, but with mean fitness $\overline{\Pi} + r$. So, it is always possible to find a value of $r$ that prevents population decline. Because this work focuses on the role of interactions, we will assume that this eigenvalue is negative and does not contribute to the destabilization of the community. Note, however, that communities enriched in non-cooperative interactions will require higher basal growth rates to thrive.

Knowing that $u = \mathbb{1}$ is an eigenvector of 3.1.3 and by defining the row vector $v = \Big[x_1^{SS}\Big(\overline{\Pi} - \frac{1}{N} - \frac{1}{N}\sum_{D=2}^{\infty}\binom{N-2}{D-2}/N^{D-2}\mathbb{E}\big[\langle {}_D\alpha_{\text{-};\text{-}}\rangle\big]\Big), x_2^{SS}\Big(\overline{\Pi} - \frac{1}{N} - \frac{1}{N}\sum_{D=2}^{\infty}\binom{N-2}{D-2}/N^{D-2}\mathbb{E}\big[\langle {}_D\alpha_{\text{-};\text{-}}\rangle\big]\Big), \ldots, x_N^{SS}\Big(\overline{\Pi} - \frac{1}{N} - \frac{1}{N}\sum_{D=2}^{\infty}\binom{N-2}{D-2}/N^{D-2}\mathbb{E}\big[\langle {}_D\alpha_{\text{-};\text{-}}\rangle\big]\Big)\Big]$ with $\mathbb{E}\big[\langle {}_D\alpha_{\text{-};\text{-}}\rangle\big]$ as defined in subsection 4.1, it can be proven (see Ding and Zhou [2007]) that the eigenvalues of $J^T$ and $A\big(x^{SS}\big) = J^T + uv$ are the same except for the one associated with the eigenvector $u$, $\lambda_u = -\overline{\Pi}$, which becomes $vu + \lambda_u = -\frac{1}{N} - \frac{1}{N}\sum_{D=2}^{\infty}\binom{N-2}{D-2}/N^{D-2}\mathbb{E}\big[\langle {}_D\alpha_{\text{-};\text{-}}\rangle\big]$. The matrix $A\big(x^{SS}\big)$ has the following explicit expression:

$$A\big(x^{SS}\big)_{i,i} = x_i^{SS}\Bigg[-1 + x_i^{SS} - \frac{1}{N} - \frac{1}{N}\sum_{D=2}^{\infty}\binom{N-2}{D-2}/N^{D-2}\mathbb{E}\big[\langle {}_D\alpha_{\text{-};\text{-}}\rangle\big] - \sum_{k\neq i}^{N-1} x_k^{SS} \sum_{D=2}^{\infty} \sum^{\binom{N-2}{D-2}} {}_D\alpha_{k;i,k_2,\ldots,k_{D-2}} x_{k_1}^{SS} x_{k_2}^{SS} \ldots x_{k_{D-2}}^{SS}\Bigg]$$

$$A\big(x^{SS}\big)_{i,j} = x_j^{SS}\Bigg[x_i^{SS} - \frac{1}{N} - \frac{1}{N}\sum_{D=2}^{\infty}\binom{N-2}{D-2}/N^{D-2}\mathbb{E}\big[\langle {}_D\alpha_{\text{-};\text{-}}\rangle\big] + \sum_{D=2}^{\infty} \sum^{\binom{N-2}{D-2}} \Big({}_D\alpha_{j;i,k_2,\ldots,k_{D-2}} x_{k_1}^{SS} x_{k_2}^{SS} \ldots x_{k_{D-2}}^{SS} - \sum_{k\neq i}^{N-1} x_k^{SS}\, {}_D\alpha_{k;i,k_2,\ldots,k_{D-2}} x_{k_1}^{SS} x_{k_2}^{SS} \ldots x_{k_{D-2}}^{SS}\Big)\Bigg] \quad (3.1.4)$$

To the best of our knowledge, there is no way to deal with the calculation of the eigenvalues of $A\big(x^{SS}\big)$, even under the assumptions considered until now. However, due to the fact that in all of the simulations checked the values $x_i^{SS}$ are or the order of $1/N$, and given the conclusions of the previous section (essentially that, in networks dominated by high-order interactions, increasing $N$ allows the existence of a stationary state arbitrarily close to being homogeneous), we hypothesized that the eigenvalues of $A\big(x^{SS}\big)$ evaluated in $x_i^{ss} = 1/N$ could be used as a reference to estimate the stability of the system.

The matrix $A\big(\frac{1}{N}\big)$ has the following form:

$$A\Big(\frac{1}{N}\Big)_{i,i} = -\frac{1}{N}\Bigg(1+\sum_{D=2}^{\infty}\binom{N-2}{D-2}/N^{D-2}\mathbb{E}\big[\langle_D\alpha_{\text{-};\text{-}}\rangle\big]\Bigg]+$$

$$+\Big[1-1/N\Big]\Bigg[\sum_{D=2}^{\infty}\binom{N-2}{D-2}/N^{D-2}\big(\langle_D\alpha_{\text{-};i}\rangle-\mathbb{E}\big[\langle_D\alpha_{\text{-};i}\rangle\big]\big)\Bigg]\Bigg) \tag{3.1.5}$$

$$A\Big(\frac{1}{N}\Big)_{i,j} = \frac{1}{N}\Bigg(\sum_{D=2}^{\infty}\binom{N-2}{D-2}/N^{D-2}\Big(\langle_D\alpha_{j;i,\text{-}}\rangle-\mathbb{E}\big[\langle_D\alpha_{j;i}\rangle\big]\Big)+$$

$$+\Big[1-1/N\Big]\Bigg[\sum_{D=2}^{\infty}\binom{N-2}{D-2}/N^{D-2}\Big(\langle_D\alpha_{\text{-};i}\rangle-\mathbb{E}\big[\langle_D\alpha_{\text{-};i}\rangle\big]\Big)\Bigg]\Bigg)$$

where $\mathbb{E}[\,\cdot\,]$ denotes the expected value, $\langle_D\alpha_{\text{-};i,\text{-}}\rangle = \frac{1}{N-1}\sum_{k\neq i}^{N-1}\langle_D\alpha_{k;i,\text{-}}\rangle$ represents the mean effect of species $i$ over the rest through order $D$ interactions; $\langle_D\alpha_{\text{-};\text{-}}\rangle = \frac{1}{N}\sum_i^N\sum_{k_l\neq i}^{\binom{N-1}{D-1}}\alpha_{i;k_1,\ldots,k_{D-1}}/\binom{N-1}{D-1}$ is the the mean impact of all order $D$ interactions, excluding the self-inhibition term; and $\langle_D\alpha_{j;i\text{-}}\rangle$ is the mean effect of species $i$ over species $j$ through order $D$ interactions. (To derive this expression, we used the equalities in 4.1.9.)

It follows from 3.1.5 that $A(\frac{1}{N})$ can be understood as a random matrix, whose entries are drawn from zero-mean distributions. Because of this, its spectrum can be derived in a completely analogous way to the matrix $I + L^{-1}\Delta$ (see subsection 4.5). Thus, assuming that the perturbation $\Big[1-1/N\Big]\Big[\sum_{D=2}^{\infty}\binom{N-2}{D-2}/N^{D-2}\big(\langle_D\alpha_{\text{-};i}\rangle-\mathbb{E}\big[\langle_D\alpha_{\text{-};i}\rangle\big]\big)\Big]$ of the otherwise constant diagonal term does not affect the distribution of the eigenvalues (see subsection 4.5 for discussion in that regard), the real parts of the eigenvalues of $A(\frac{1}{N})$ are bounded, in the limit $N\to\infty$, by the following expression:

$$Re\big[\lambda_{A(\frac{1}{N})}\big] \leq \Big(1+\rho\Big[A(\tfrac{1}{N})_{i,j},A(\tfrac{1}{N})_{j,i}\Big]\Big)\sigma\Big(A(\tfrac{1}{N})_{i,j}\Big)N^{1/2}-1-\sum_{D=2}^{\infty}\binom{N-2}{D-2}/N^{D-2}\mathbb{E}\big[\langle_D\alpha_{\text{-};\text{-}}\rangle\big] \tag{3.1.6}$$

where $Re\big[\lambda_{A(\frac{1}{N})}\big]$ indicates the real part of the eigenvalues of $A(\frac{1}{N})$, and $\sigma$ and $\rho$ denote the standard deviation and correlation coefficient, respectively. Given the values of $\mathbb{E}\big[\langle_D\alpha_{\text{-};\text{-}}\rangle\big]$ (see 4.1.5), the variance of $\langle_D\alpha_{i;j,\text{-}}\rangle$ (see 4.1.7), and the covariance of $\langle_D\alpha_{i;j,\text{-}}\rangle$ and $\langle_D\alpha_{j;i,\text{-}}\rangle$ (see 4.1.14); inequality 3.1.6 becomes:

$$\begin{gathered}
Re\big[\lambda_{A(\frac{1}{N})}\big] \leq (1+\rho)\sigma-1-\mu \\
\text{with } \mu = \sum_{D=2}^{\infty}\binom{N-2}{D-2}/N^{D-2}\phi_D C_D\big(2P_D-1\big) \\
\text{and } \sigma = \Bigg[\sum_{D=2}^{\infty}\binom{N-2}{D-2}\frac{\phi_D^2 C_D}{N^{2D-5}}\Big[1-C_D\big(2P_D-1\big)^2\Big]\Bigg]^{1/2} \\
\text{and } \rho = \frac{\sum_{D=2}^{\infty}\binom{N-2}{D-2}\frac{\phi_D^2 C_D}{N^{2(D-2)}}\Big(\Big[2S_D-1-C_D\big(2P_D-1\big)^2\Big]-A_D\Big[2S_D-1\Big]\Big)}{\sum_{D=2}^{\infty}\binom{N-2}{D-2}\frac{\phi_D^2 C_D}{N^{2(D-2)}}\Big[1-C_D\big(2P_D-1\big)^2\Big]}
\end{gathered} \tag{3.1.7}$$

where the impact of terms $\big[1-1/N\big]\Big[\sum_{D=2}^{\infty}\binom{N-2}{D-2}/N^{D-2}\big(\langle_D\alpha_{\text{-};i}\rangle-\mathbb{E}\big[\langle_D\alpha_{\text{-};i}\rangle\big]\big)\Big]$ has been neglected in the calculation of the variance and $\rho$. In the limit $N\to\infty$:

$$
\begin{aligned}
\lim_{N\to\infty} Re\big[\lambda_{A(\frac{1}{N})}\big] &\leq \big(1+\rho\big)\sigma - 1 - \mu \\
\text{with } \mu &= \sum_{D=2}^{\infty} \frac{\phi_D C_D\big(2P_D - 1\big)}{(D-2)!} \\
\text{and } \sigma &= \left[\sum_{D=2}^{\infty} \frac{\phi_D^2 C_D}{(D-2)!}\left[1 - C_D\big(2P_D - 1\big)^2\right] N^{3-D}\right]^{1/2} \\
\text{and } \rho &= \frac{\sum_{D=2}^{\infty} \frac{\phi_D^2 C_D}{(D-2)!}\left(\left[2S_D - 1 - C_D\big(2P_D-1\big)^2\right] - A_D\left[2S_D - 1\right]\right) N^{2-D}}{\sum_{D=2}^{\infty} \frac{\phi_D^2 C_D}{(D-2)!}\left[1 - C_D\big(2P_D-1\big)^2\right] N^{2-D}}
\end{aligned}
\tag{3.1.8}
$$

From the expressions above, an approximate analytical condition for the stability of the stationary state results from imposing that the right-hand-side of the inequality in 3.1.7 is smaller than zero (hence, the largest eigenvalue $Re\big[\lambda_{A(\frac{1}{N})}\big]$ necessarily becomes negative). In that way, we obtain the expression $1 + \mu > \sigma(1 + \rho)$, which corresponds to eqn. 2 in the main text.

#### 3.2 Numerical validation

To check to which extent expression 3.1.7 can be used as an approximate stability criterion, simulations were performed covering a wide range of conditions. As main variables, we focused on the number of species $N$ and the interaction diversity (or more precisely, the diversity of interaction profiles). For single order networks, the latter is defined as the fraction of all possible interaction profiles in terms of connectivity, positivity and symmetry that are stable given a number of species $N$ and an interaction strength. For networks with interactions of different orders, this idea can be easily generalised by considering the connectivities, positivities and symmetries of the orders included, along with the relative interaction strengths of these orders (note that by dividing the interaction strength associated with a particular order by the maximum strength of all orders, these relative strengths are bounded between zero and one).

Apart from the number of species $N$ and the interaction diversity, five specific cases were explored: three associated with networks composed just by single order interactions (orders 2,3 and 4); and two mixed interaction networks involving a mixture of order 2,3 and 4 interactions where the relative interaction strengths are normalized in different ways (mean and variance normalization). The mean normalization sets the relative interaction strengths in a way that, if two orders share the same values of connectivity, positivity and symmetry, the $\mu$ term corresponding to both orders in condition 3.1.7 takes the same value. The variance normalization sets the relative interaction strengths in a way that, if two orders share the same values of connectivity, positivity and symmetry, the $\sigma$ term corresponding to both orders in condition 3.1.7 takes the same value. Note that these two normalization approaches correspond to two very different scenarios: the mean normalization makes the impact of the order-2 interactions negligible in the $\sigma$ term when $N$ is high enough. The variance normalization, on the other hand, allows a comparable contribution of different interaction orders in the $\sigma$ term at the expense of order 4 interactions being dominant in the $\mu$ term.

For each of these five cases, a total of 9604 stable and 9604 unstable networks according to the approximate stability criterion 3.1.7 were randomly selected for several values of $N$ and interaction diversity. The community dynamics of these networks were then simulated (see Methods: Numerical simulations of community dynamics in the main text) to check how many did not match their predicted stability (see Figure 1). Because the approximate stability criterion assumes the existence of a stationary state, it is possible that a network is correctly predicted to be stable although the corresponding simulation fails to reach a stationary state. Thus, only those networks of the 9604 sampled for which a steady state was found *via* Newton-Raphson method (starting point: homogeneous state) were considered in the analysis. If less than 1000 networks fulfilling this condition were found, the corresponding error rate is not displayed (gray areas in Fig. 1). The number of simulations was chosen to allow estimation of proportions with a 95% confidence interval narrower than 0.01 under the normal approach.

The results of the simulations are clear: the approximate stability criterion 3.1.7 is a good reference for understanding the effect of the different network parameters on the stability of the system. Moreover, the error diminishes as $N$ and the interaction diversity increase, for the stable region, and as $N$ increases and the interaction diversity decreases, for the unstable region.

### 4 Auxiliary results.

#### 4.1 Some properties of the $\langle_D \alpha_{i;j,\text{-}}\rangle$, $\langle_D \alpha_{i;\text{-}}\rangle$, $\langle_D \alpha_{\text{-};j,\text{-}}\rangle$ and $\langle_D \alpha_{\text{-};\text{-}}\rangle$ distributions:

Under the assumptions about the interaction network stated in *Section 1*, the term $\langle_D \alpha_{i;j,\text{-}}\rangle$ can be formulated as follows:

$$\langle_D \alpha_{i;j,\text{-}}\rangle = \frac{\phi_D(n_+ - n-)}{\binom{N-2}{D-2}} \tag{4.1.1}$$

where $\phi_D$ indicates the strength of order $D$ interactions and $n_+$ ($n_-$) represents the number of interactions involving $i$ and $j$ in which $i$ is positively (negatively) affected. Under the assumption of unstructured networks, the variables $n_+$ and $n_-$, along with $n_0$ (the number of interactions involving $i$ and $j$ in which $i$ is unaffected) follows a multinomial distribution with parameters $C_D P_D$, $C_D(1-P_D)$ and $(1-C_D)$, where $C_D$ is the probability of an species being affected (either in a positive or in a negative way) by an interaction:

$$C_D = \sum_{k_+=0}^{D} \sum_{k_-=0}^{D-k_+} \frac{k_+ + k_-}{D} p_{(k_+,k_-)} \tag{4.1.2}$$

and $P_D$ is the positivity of the order $D$ interactions, defined as the conditional probability of a species benefiting from an interaction given that it is affected by such interaction:

$$P_D = \left( \sum_{k_+=0}^{D} \sum_{k_-=0}^{D-k_+} \frac{k_+}{D} p_{(k_+,k_-)} \right) \Big/ C_D \tag{4.1.3}$$

where $p_{(k_+,k_-)}$ represents the probability of an interaction affecting $k_+$ species in a positive way and $k_-$ species in a negative way while the $D - k_+ - k_-$ remaining species are unaffected.

Given this parametrization and the properties of the mean:

$$\mathbb{E}\big[\langle_D \alpha_{i;j,\text{-}}\rangle\big] = \frac{\phi_D}{\binom{N-2}{D-2}} \Big(\mathbb{E}[n_+] - \mathbb{E}[n-]\Big) \tag{4.1.4}$$

where $\mathbb{E}[\cdot]$ indicates the expected value of the corresponding random variable. According to the properties of multinomial distributions $\mathbb{E}[n_+] = \binom{N-2}{D-2} C_D P_D$ and $\mathbb{E}[n_-] = \binom{N-2}{D-2} C_D \big(1 - P_D\big)$ so:

$$\mathbb{E}\big[\langle_D \alpha_{i;j,\text{-}}\rangle\big] = \phi_D C_D \Big(2P_D - 1\Big) \tag{4.1.5}$$

Analogously, given the properties of the variance:

$$\sigma^2_{\langle_D \alpha_{i;j,\text{-}}\rangle} = \frac{\phi_D^2}{\binom{N-2}{D-2}^2}\Big(\sigma^2_{n_+} + \sigma^2_{n-} - 2cov(n_+, n-)\Big) \tag{4.1.6}$$

where $\sigma^2$ denotes the variance and $cov(x, y)$ the covariance of $x$ and $y$. Substituting the values of the multinomial distribution and doing some algebra:

$$\sigma^2_{\langle_D \alpha_{i;j,\text{-}}\rangle} = \frac{\phi_D^2 C_D}{\binom{N-2}{D-2}}\Big[1 - C_D\big(2P_D - 1\big)^2\Big] \tag{4.1.7}$$

The variance of $\langle_D \alpha_{i;\text{-}}\rangle = \frac{1}{\binom{N-1}{D-1}} \sum_{k_l \neq i}^{\binom{N-1}{D-1}} \alpha_{i;k_1,\ldots,k_{D-1}}$ can be calculated in a completely analogous way, yielding:

$$\sigma^2_{\langle_D \alpha_{i;\text{-}}\rangle} = \frac{\phi_D^2 C_D}{\binom{N-1}{D-1}}\Big[1 - C_D\big(2P_D - 1\big)^2\Big] \tag{4.1.8}$$

Because of the assumption of random interaction networks, all $\langle_D \alpha_{i;j,\text{-}}\rangle$ follow the same distribution and thus, have the same properties. Knowing the expected value of $\langle_D \alpha_{i;j,\text{-}}\rangle$, the corresponding values for distributions $\langle_D \alpha_{i;\text{-}}\rangle$, $\langle_D \alpha_{\text{-};j,\text{-}}\rangle = \frac{1}{N-1}\sum_{k\neq i}^{N-1}\langle_D \alpha_{k;j,\text{-}}\rangle$ and $\langle_D \alpha_{\text{-};\text{-}}\rangle = \frac{1}{N}\sum_{k}^{N}\langle_D \alpha_{k;\text{-}}\rangle$ can be readily obtained by applying the properties of the expected value:

$$\mathbb{E}\big[\langle_D \alpha_{i;j,\text{-}}\rangle\big] = \mathbb{E}\big[\langle_D \alpha_{\text{-};j,\text{-}}\rangle\big] = \mathbb{E}\big[\langle_D \alpha_{i;\text{-}}\rangle\big] = \mathbb{E}\big[\langle_D \alpha_{\text{-};\text{-}}\rangle\big] \tag{4.1.9}$$

It is also important to note that variables $\langle_D \alpha_{i;j,\text{-}}\rangle$ and $\langle_D \alpha_{k;l,\text{-}}\rangle$ may be correlated (especially if some of the indices are shared) due to the fact that interactions involving species $i, j, k, l$ affect both variables.

In addition to the variance and the expected value of $\langle_D \alpha_{i;j,\text{-}}\rangle$, the covariance of random variables $\langle_D \alpha_{i;j,\text{-}}\rangle$ and $\langle_D \alpha_{j;i,\text{-}}\rangle$ is required for estimating the distribution of eigenvalues in several occasions. To obtain this covariance, let us define $p_=$ and $p_{\neq}$ as the probabilities of two species participating in the same interaction being affected in the same way and in different ways, respectively, and $p_{*0} = 1 - p_= - p_{\neq}$ as the probability that at least one species remains unaffected. With these definitions it is clear that:

$$\begin{aligned}\mathbb{E}\big[\langle_D \alpha_{i;j,\text{-}}\rangle\langle_D \alpha_{j;i,\text{-}}\rangle\big] &= \phi_D^2\Big(p_= - p_{\neq}\Big) = \phi_D^2\Big(2p_= - 1 + p_{*0}\Big) = \\ &= \phi_D^2\big(1 - p_{*0}\big)\Big(2\frac{p_=}{1 - p_{*0}} - 1\Big) = \phi_D^2\big(1 - p_{*0}\big)\Big(2S_D - 1\Big)\end{aligned} \tag{4.1.10}$$

where the parameter $S_D$ has been defined as a measure of the degree of symmetry of the interaction network. As it can be inferred from equation 4.1.10, $S_D$ it is the probability of two species being affected in the same way by an interaction conditioned to both species being affected by such interaction. Under the model assumptions $p_=$,$p_{\neq}$ and $p_{*0}$ are independent of the species involved in the interaction so:

$$p_{=} = \sum_{k_+=0}^{D} \sum_{k_-=0}^{D-k_+} \frac{\binom{k_+}{2} + \binom{k_-}{2}}{\binom{D}{2}} p_{(k_+,k_-)} \tag{4.1.11}$$

$$\begin{aligned} p_{*0} &= \sum_{k_+=0}^{D} \sum_{k_-=0}^{D-k_+} \frac{\binom{D-k_--k_+}{2} + 2k_+(D-k_--k_+) + 2k_-(D-k_--k_+)}{\binom{D}{2}} p_{(k_+,k_-)} = \\ &= \sum_{k_+=0}^{D} \sum_{k_-=0}^{D-k_+} \frac{(D-k_--k_+)}{D} p_{(k_+,k_-)} + \sum_{k_+=0}^{D} \sum_{k_-=0}^{D-k_+} \frac{(D-k_--k_+)(k_-+k_+)}{D(D-1)} p_{(k_+,k_-)} = \\ &= 1 - C_D + A_D \end{aligned} \tag{4.1.12}$$

where the parameter $A_D \leq C_D$ has been introduced as a measure of the level of *mensalism* in the network. As equation 4.1.12 indicates, $A_D$ is the probability of one species of a randomly taken pair being unaffected by an order $D$ interaction. Given equations 4.1.12 and 4.1.10:

$$\mathbb{E}\big[\langle_D \alpha_{i;j,\text{-}}\rangle\langle_D \alpha_{j;i,\text{-}}\rangle\big] = \phi_D^2 (C_D - A_D)\Big(2S_{_D} - 1\Big) \tag{4.1.13}$$

Finally, by the definition of covariance, and knowing that under the assumptions of the model $\mathbb{E}\big[\langle_D \alpha_{i;j,\text{-}}\rangle\big] = \mathbb{E}\big[\langle_D \alpha_{i;j,\text{-}}\rangle\big]$:

$$cov\big[\langle_D \alpha_{i;j,\text{-}}\rangle, \langle_D \alpha_{j;i,\text{-}}\rangle\big] = \phi_D^2 C_D\Big[2S_{_D} - 1 - C_D\big(2P_D - 1\big)^2\Big] - \phi_D^2 A_D\Big[2S_{_D} - 1\Big] \tag{4.1.14}$$

In the main text, the analysis and discussion focus on interaction networks in which all or none of the species are affected by the interaction (the latter case is simply interpreted as an absence of interaction). In those cases $A_D = 0$ for all $D$ and $C_D$ is just the fraction of realized interactions in the order $D$ network.

#### 4.2 Upper bound for the maximum of a sample coming from a half-mean distribution:

The expectation of the maximum of $n$ draws of a random variable can be bounded using the corresponding moment generating function ($M_X(t)$) by using the following procedure from J.G [2019]:

$$e^{t\big\langle \max(X_1,X_2,...,X_n)\big\rangle} \leq \big\langle e^{t \;\max(X_1,X_2,...,X_n)}\big\rangle \leq \big\langle \sum_{i=1}^{n} e^{tX_i}\big\rangle = nM_X(t) \tag{4.2.1}$$

where the first inequality is due to Jensen's inequality. Taking logarithms:

$$\big\langle \max(X_1, X_2, ..., X_n)\big\rangle \leq \frac{log(n)}{t} + \frac{log(M_X(t))}{t} \tag{4.2.2}$$

Because this inequality holds for every $t$, the best bound is obtained by minimizing the right side expression. The moment generating function of a half-normal random variable $X$ is:

$$M_X(t) = 2\Phi(\sigma t)e^{\frac{\sigma^2 t^2}{2}} \leq 2e^{\frac{\sigma^2 t^2}{2}} \tag{4.2.3}$$

where $\Phi$ represents the cumulative distribution function of the standard normal; and $\sigma$ is the standard deviation of the normal distribution associated to $X$.

Considering inequalities 4.2.2 and 4.2.3, the following upper bound for $\big\langle \max(X_1, X_2, ..., X_n)\big\rangle$ is obtained choosing $t = \frac{\sqrt{2log(2n)}}{\sigma}$:

$$\big\langle \max(X_1, X_2, ..., X_n)\big\rangle \leq \sqrt{2log(n)}\sigma \tag{4.2.4}$$

#### 4.3 Upper bound for $\big|L^{-1}F(x_0)\big|_\infty$:

By definition of the infinity norm for vectors:

$$\big|L^{-1}F(x_0)\big|_\infty = \frac{1}{|\beta|}\max_i\left(\Bigg|\sum_{D=2}^{\infty}\binom{N-1}{D-1}/N^{D-1}\Big[\langle_D\alpha_{i;\text{-}}\rangle - \langle_D\alpha_{\text{-};\text{-}}\rangle\Big]\Bigg|\right)$$
$$\text{with } \beta = 1 + \sum_{D=2}^{\infty}\binom{N-2}{D-2}/N^{D-2}\mathbb{E}\big[\langle_D\alpha_{\text{-};\text{-}}\rangle\big] \tag{4.3.1}$$

Due to the random nature of the network, the value $\max_i\left(\Big|\sum_{D=2}^{\infty}\binom{N-1}{D-1}/N^{D-1}\Big[\langle_D\alpha_{i;\text{-}}\rangle - \langle_D\alpha_{\text{-};\text{-}}\rangle\Big]\Big|\right)$ follows a probability distribution. Thus, the value of its expected value $\mathbb{E}\left[\left(\Big|\sum_{D=2}^{\infty}\binom{N-1}{D-1}/N^{D-1}\Big[\langle_D\alpha_{i;\text{-}}\rangle - \langle_D\alpha_{\text{-};\text{-}}\rangle\Big]\Big|\right)\right]$ will be used as a reference. To obtain an upper bound of this expectation, first of all, it is important to consider that the terms $\langle_D\alpha_{i;\text{-}}\rangle - \langle_D\alpha_{\text{-};\text{-}}\rangle = \langle_D\alpha_{i;\text{-}}\rangle - \mathbb{E}\big[\langle_D\alpha_{i;\text{-}}\rangle\big] + \mathbb{E}\big[\langle_D\alpha_{\text{-};\text{-}}\rangle\big] - \langle_D\alpha_{\text{-};\text{-}}\rangle \sim \langle_D\alpha_{i;\text{-}}\rangle - \mathbb{E}\big[\langle_D\alpha_{i;\text{-}}\rangle\big]$ closely follow a normal distribution with mean zero and variance $\sigma^2_{\langle_D\alpha_{i;\text{-}}\rangle} = \frac{\phi_D^2 C_D}{\binom{N-1}{D-1}}\Big[1 - C_D\big(2P_D - 1\big)^2\Big]$ (see equation 4.1.8). This is due to the fact that each term comes from the sum of $\binom{N-1}{D-1}$ independent (and well behaved) random variables and thus, if $N$ is big enough, the Central Limit Theorem can be applied. Under this approximation, given the properties of the normal distribution, and due to the assumption of different orders being independent; the term $\sum_{D=2}^{\infty}\binom{N-1}{D-1}/N^{D-1}\Big[\langle_D\alpha_{i;\text{-}}\rangle - \langle_D\alpha_{\text{-};\text{-}}\rangle\Big]$ can be approached to a normal distribution of mean zero and variance $\sigma^2_{L^{-1}F(x_0)} = \sum_{D=2}^{\infty}\binom{N-1}{D-1}^2/N^{2(D-1)}\sigma^2_{\langle_D\alpha_{i;\text{-}}\rangle}$ when $N$ is big.

But if the distribution of $\sum_{D=2}^{\infty}\binom{N-1}{D-1}/N^{D-1}\Big[\langle_D\alpha_{i;\text{-}}\rangle - \langle_D\alpha_{\text{-};\text{-}}\rangle\Big]$ can be approached to the previously specified zero-mean normal, then its absolute value follows the corresponding half-normal distribution. Knowing this, the expectation of the maximum of $N$ draws of this random variable can be bounded using the corresponding moment generating function as described in subsection 4.2. Finally, combining expressions 4.3.1, 4.2.4, 4.1.9, 4.1.5 and the value of $\sigma^2_{L^{-1}F(x_0)}$ obtained above:

$$\begin{aligned}\left|L^{-1}F(x_0)\right|_\infty \leq \frac{\sqrt{2log(N)}}{|\beta|}\left(\sum_{D=2}^{\infty}\binom{N-1}{D-1}/N^{2(D-1)}\phi_D^2 C_D\left[1-C_D\left(2P_D-1\right)^2\right]\right)^{1/2}\\ \text{with } \beta = 1+\sum_{D=2}^{\infty}\phi_D C_D\left(2P_D-1\right)\binom{N-2}{D-2}/N^{D-2}\end{aligned} \tag{4.3.2}$$

that when $N \to \infty$:

$$\begin{aligned}\lim_{N\to\infty}\left|L^{-1}F(x_0)\right|_\infty \leq \frac{\sqrt{2log(N)}}{|\beta|}\left(\sum_{D=2}^{\infty}N^{1-D}\frac{\phi_D^2 C_D}{(D-1)!}\left[1-C_D\left(2P_D-1\right)^2\right]\right)^{1/2}\\ \text{with } \beta = 1+\sum_{D=2}^{\infty}\frac{\phi_D C_D\left(2P_D-1\right)}{(D-2)!}\end{aligned} \tag{4.3.3}$$

#### 4.4 Upper and lower bounds for $\left|\left(I + L^{-1}\Delta\right)^{-1}\right|_\infty$:

Only the case of $I + L^{-1}\Delta$ being strictly diagonally dominant (*i.e.* the absolute value of the $i^{th}$ diagonal element is greater than the sum of the absolute values of the rest of the $i^{th}$ row, $|a_{i,i}| > \sum_{j\neq i}|a_{i,j}|$) will be discussed in relation with the calculation of the upper bound. In the remaining scenarios, or when the bound is not applicable or tight enough, the exact value of the norm is obtained numerically.

Under the above assumption, the Varah bound (Varah [1975]) can be applied:

$$\left|A^{-1}\right|_\infty \leq \frac{1}{\min_i\left(|a_{i,i}| - \sum_{j\neq i}|a_{i,j}|\right)} \tag{4.4.1}$$

being $A$ any strictly diagonally dominant matrix and $a_{i,j}$ the corresponding matrix element. Let us define $L^{-1}\Delta[i,j]$ as the $i,j$ term of matrix $L^{-1}\Delta$. Then, by applying inequality 4.4.1 and some minor considerations:

$$\left|\left(I + L^{-1}\Delta\right)^{-1}\right|_\infty \leq \frac{1}{\min_i\left(|1+L^{-1}\Delta[i,i]| + |L^{-1}\Delta[i,i]| - \sum_j |L^{-1}\Delta[i,j]|\right)} \leq \frac{1}{1-|L^{-1}\Delta|_\infty} \tag{4.4.2}$$

Because the infinity norm is the maximum absolute row sum of the matrix, $\left|L^{-1}\Delta\right|_\infty$ can be trivially bounded by the maximum absolute value of the whole matrix $L^{-1}\Delta$, $max\left(|L^{-1}\Delta|\right)$. Considering this and equation 2.2.5:

$$\left|L^{-1}\Delta\right|_{\infty} \leq N max\Big(|L^{-1}\Delta|\Big) = \frac{N}{|\beta|} max\Big(|\Delta_{i,j}|\Big) \sim \frac{N}{|\beta|} max\left(\left|\sum_{D=2}^{\infty}\binom{N-2}{D-2}/N^{D-2}\Big[\langle_D\alpha_{i;j,\text{-}}\rangle - \mathbb{E}\big[\langle_D\alpha_{\text{-};\text{-}}\rangle\big]\Big]\right|\right)$$
$$\text{with } \beta = 1+\sum_{D=2}^{\infty}\binom{N-2}{D-2}/N^{D-2}\mathbb{E}\big[\langle_D\alpha_{\text{-};\text{-}}\rangle\big] \tag{4.4.3}$$

If $N$ is big enough, the contribution of the term $\left(1-\frac{1}{N}\right)\binom{N-2}{D-2}/N^{D-2}\Big[\langle_D\alpha_{\text{-};i,\text{-}}\rangle - \mathbb{E}\big[\langle_D\alpha_{\text{-};\text{-}}\rangle\big]\Big]$ in the expression of $\Delta_{i,j}$ (equation 2.2.5) can be neglected, and so it was not considered in the inequality 4.4.3. Moreover, the diagonal terms of $L^{-1}\Delta$ are almost always smaller than those of the off-diagonals, so only the cases $i \neq j$ can be considered in the estimation of $\max_{i,j}\Big(|\Delta_{i,j}|\Big)$.

Due to the random nature of the network, the value $\max_{i,j}\Big(|\Delta_{i,j}|\Big)$ follows a probability distribution. Thus, the value of its mean $\mathbb{E}\Big[\max_{i,j}\Big(|\Delta_{i,j}|\Big)\Big]$ will be used as a reference. To obtain an upper bound of this expectation, first of all, it is important to consider that, for orders greater than 2, the terms $\langle_D\alpha_{i;j,\text{-}}\rangle - \langle_D\alpha_{\text{-};\text{-}}\rangle$ closely follow a normal distribution with mean zero and variance that of equation 4.1.7. This is due to the fact that each term comes from the sum of $\binom{N-2}{D-2}$ independent (and well behaved) random variables and thus, if $N$ is big enough, the Central Limit Theorem can be applied. Under this approximation, given the properties of the normal distribution, and due to the assumption of different orders being independent; the term $\sum_{D=3}^{\infty}\binom{N-2}{D-2}/N^{D-2}\Big[\langle_D\alpha_{i,;j,\text{-}}\rangle - \mathbb{E}\big[\langle_D\alpha_{\text{-};\text{-}}\rangle\big]\Big]$ can be approached to a normal distribution of mean zero and variance $\sum_{D=3}^{\infty}\binom{N-2}{D-2}^2/N^{2D-4}\ \sigma^2_{\langle_D\alpha_{i,;j,\text{-}}\rangle}$ when $N$ is big.

Unfortunately, the same reasoning cannot be applied to order 2 interactions because, in this case, species $i$ is affected by species $j$ only through one interaction. However, due to the fact that $\langle_2\alpha_{i,;j,\text{-}}\rangle$ can only take three values ($\phi_2$,$-\phi_2$ or 0), and given the high number of elements in matrix $L^{-1}\Delta$, it is reasonable to establish an upper bound of $\mathbb{E}\Big[\max_{i,j}\Big(|\Delta_{i,j}|\Big)\Big]$ as follows:

$$\begin{aligned}
&\mathbb{E}\Big[\max_{i,j}\Big(|\Delta_{i,j}|\Big)\Big] \sim \mathbb{E}\left[\max_{i,j}\left(\left|\sum_{D=2}^{\infty}\binom{N-2}{D-2}/N^{D-2}\Big[\langle_D\alpha_{i;j,\text{-}}\rangle - \mathbb{E}\big[\langle_D\alpha_{\text{-};\text{-}}\rangle\big]\Big]\right|\right)\right] \leq \\
&\leq \mathbb{E}\left[\max_{i,j}\left(\Big|\langle_2\alpha_{i;j}\rangle - \mathbb{E}\big[\langle_2\alpha_{\text{-};\text{-}}\rangle\big]\Big| + \left|\sum_{D=3}^{\infty}\binom{N-2}{D-2}/N^{D-2}\Big[\langle_D\alpha_{i;j,\text{-}}\rangle - \mathbb{E}\big[\langle_D\alpha_{\text{-};\text{-}}\rangle\big]\Big]\right|\right)\right] \leq \\
&\leq \phi_2\big(1+C_2|2P_2-1|\big) + \mathbb{E}\left[\max_{i,j}\left(\left|\sum_{D=3}^{\infty}\binom{N-2}{D-2}/N^{D-2}\Big[\langle_D\alpha_{i;j,\text{-}}\rangle - \mathbb{E}\big[\langle_D\alpha_{\text{-};\text{-}}\rangle\big]\Big]\right|\right)\right]
\end{aligned} \tag{4.4.4}$$

where $\mathbb{E}\big[\langle_2\alpha_{\text{-};\text{-}}\rangle\big] = \phi_2C_2\big(2P_2-1\big)$ (see 4.1.5) was used for obtaining the last inequality. But if the distribution of $\sum_{D=3}^{\infty}\binom{N-2}{D-2}/N^{D-2}\Big[\langle_D\alpha_{i,;j,\text{-}}\rangle - \mathbb{E}\big[\langle_D\alpha_{\text{-};\text{-}}\rangle\big]\Big]$ can be approximated by the previously specified zero-mean normal, then its absolute value follows the corresponding half-normal distribution. Knowing this, and substituting expression 4.2.4 from subsection 4.2 with $n = N^2$ (the number of components of the matrix $L^{-1}\Delta$) in inequality 4.4.4, and then the corresponding result in in 4.4.3; considering expressions 4.1.5 and 4.1.7; and knowing that $\sigma^2 = \sum_{D=3}^{\infty}\binom{N-2}{D-2}^2/N^{2D-4}\ \sigma^2_{\langle_D\alpha_{i,;j,\text{-}}\rangle}$, the desired upper bound is obtained:

$$\left|L^{-1}\Delta\right|_\infty \leq \frac{\phi_2\big(1+C_2|2P_2-1|\big)}{|\beta|}N + \frac{2\sqrt{log(N)}}{|\beta|}\left[\sum_{D=3}^{\infty}\binom{N-2}{D-2}/N^{2(D-3)}\phi_D^2 C_D\Big(1-C_D\Big[1-4P_D(1-P_D)\Big]\Big)\right]^{1/2}$$
$$\text{with } \beta = 1+\sum_{D=2}^{\infty}\phi_D C_D\big(2P_D-1\big)\binom{N-2}{D-2}/N^{D-2} \tag{4.4.5}$$

that when $N\to\infty$:

$$\lim_{N\to\infty}\left|L^{-1}\Delta\right|_\infty \leq \frac{\phi_2\big(1+C_2|2P_2-1|\big)}{|\beta|}N + \frac{2\sqrt{log(N)}}{|\beta|}\left[\sum_{D=3}^{\infty}\frac{N^{4-D}}{(D-2)!}\phi_D^2 C_D\Big(1-C_D\Big[1-4P_D(1-P_D)\Big]\Big)\right]^{1/2}$$
$$\text{with } \beta = 1+\sum_{D=2}^{\infty}\frac{\phi_D C_D}{(D-2)!}\big(2P_D-1\big) \tag{4.4.6}$$

As it is clear from equation 4.4.6, the upper bound for $\left|\left(I+L^{-1}\Delta\right)^{-1}\right|_\infty$ defined in expression 4.4.2 is only useful for interaction networks dominated by order $\geq 5$ interaction, specially for large $N$, when the value of 4.4.6 tends to 1. In the remaining scenarios, a direct numerical calculation of either $\left|L^{-1}\Delta\right|_\infty$ or $\left|\left(I+L^{-1}\Delta\right)^{-1}\right|_\infty$ is almost always required.

In those cases where the upper bound cannot be estimated, it is interesting to have an idea of how low can $\left|\left(I+L^{-1}\Delta\right)^{-1}\right|_\infty$ be. If using this lower bound the conditions for the application of the Kantorovich Theorem are not fulfilled, then they are not fulfilled for sure if the actual value of $\left|\left(I+L^{-1}\Delta\right)^{-1}\right|_\infty$ is used. To estimate this lower bound the following chain of inequalities is used:

$$\left|\left(I+L^{-1}\Delta\right)^{-1}\right|_\infty \geq r\Big(\big[I+L^{-1}\Delta\big]^{-1}\Big) = \frac{1}{\min\limits_i|\lambda_i|} \geq \frac{1}{|\lambda_i|} \tag{4.4.7}$$

where $r\Big(\big[I+L^{-1}\Delta\big]^{-1}\Big)$ is the spectral radius of $\big[I+L^{-1}\Delta\big]^{-1}$ and $\lambda_i$ represents an eigenvalue of $I+L^{-1}\Delta$. According to subsection 4.5, the eigenvalues of $I+L^{-1}\Delta$ are distributed in an ellipse centered in 1. Thus, there exist eigenvalues that fulfill $|\lambda_i|\leq 1$. Thus, according to inequalities 4.4.7:

$$\left|\left(I+L^{-1}\Delta\right)^{-1}\right|_\infty \geq 1 \tag{4.4.8}$$

In particular, and according to subsection 4.5 $|\lambda_i|\to 1$ as $N\to\infty$ if no order 2 and 3 interactions are included. In the rest of situations, increasing $N$ either does not alter significantly the spectrum of $I+L^{-1}\Delta$ (no order 2 interactions); or increases the size of the ellipse even to the point of including the point $(0,0)$ in its area, with the concomitant increase in $\left|\left(I+L^{-1}\Delta\right)^{-1}\right|_\infty$.

#### 4.5 Spectrum of $I+L^{-1}\Delta$:

Considering expression 4.1.9 in subsection 4.1, the matrix $L^{-1}\Delta$ (see equation 2.2.5) can be rewritten as:

$$L^{-1}\Delta = -1/\beta \begin{bmatrix} \Delta_{1,1} & \Delta_{1,2} & \cdots & \Delta_{1,N-1} & \Delta_{1,N} \\ \Delta_{2,1} & \Delta_{2,2} & \cdots & \Delta_{2,N-1} & \Delta_{2,N} \\ \vdots & \vdots & \ddots & \vdots & \vdots \\ \Delta_{N-1,1} & \Delta_{N-1,2} & \cdots & \Delta_{N-1,N-1} & \Delta_{N-1,N} \\ \Delta_{N,1} & \Delta_{N,2} & \cdots & \Delta_{N,N-1} & \Delta_{N,N} \end{bmatrix}$$

$$\begin{aligned} &\text{with } \beta = 1 + \sum_{D=2}^{\infty} \binom{N-2}{D-2} / N^{D-2} \mathbb{E}\big[\langle {}_D\alpha_{\text{-};\text{-}} \rangle\big] \\ &\text{and } \Delta_{i,i} = \sum_{D=2}^{\infty} -\left(1 - \frac{1}{N}\right) \binom{N-2}{D-2} / N^{D-2} \Big( \langle {}_D\alpha_{\text{-};i,\text{-}} \rangle - \mathbb{E}\big[\langle {}_D\alpha_{i;\text{-}} \rangle\big] \Big) \end{aligned} \tag{4.5.1}$$

$$\begin{aligned} \text{and } \Delta_{i,j} = &\sum_{D=2}^{\infty} \binom{N-2}{D-2} / N^{D-2} \left[ \langle {}_D\alpha_{i;j,\text{-}} \rangle - \mathbb{E}\big[\langle {}_D\alpha_{i;j,\text{-}} \rangle\big] + \Big( \langle {}_D\alpha_{\text{-};j,\text{-}} \rangle - \mathbb{E}\big[\langle {}_D\alpha_{\text{-};j,\text{-}} \rangle\big] \Big) \right] + \\ &+ \frac{1}{N} \sum_{D=2}^{\infty} \binom{N-2}{D-2} / N^{D-2} \Big( \langle {}_D\alpha_{\text{-};j,\text{-}} \rangle - \mathbb{E}\big[\langle {}_D\alpha_{\text{-};j,\text{-}} \rangle\big] \Big) \end{aligned}$$

According to equation 4.5.1, it is clear that matrix $L^{-1}\Delta$ can be understood as a random matrix whose entries are drawn from zero-mean distributions. Additionally, it is important to note that, as commented in subsection 4.1, the values of $\langle {}_D\alpha_{i;j,\text{-}} \rangle$ and $\langle {}_D\alpha_{k;l,\text{-}} \rangle$, and concomitantly the corresponding matrix entries, are expected to present a certain degree of correlation depending on the shared indexes. According to Baron et al. [2022] (where the eigenvalues of random matrices with generalised correlations are characterized), the eigenvalues of $I + L^{-1}\Delta$ are distributed in the following ellipse in the limit of $N \to \infty$:

$$\begin{aligned} \frac{(x-1)^2}{(1+\rho(\Delta_{i,j}, \Delta_{j,i}))^2} + \frac{y^2}{(1-\rho(\Delta_{i,j}, \Delta_{j,i}))^2} &= \frac{\sigma^2(\Delta_{i,j})}{(1+\beta)^2} N \\ \text{with } \beta = 1 + \sum_{D=2}^{\infty} \binom{N-2}{D-2} / N^{D-2} \mathbb{E}\big[\langle {}_D\alpha_{\text{-};\text{-}} \rangle\big] \end{aligned} \tag{4.5.2}$$

where $\sigma$ and $\rho$ indicate the standard deviation and correlation coefficient, respectively. However, expression 4.5.2 is strictly valid if the diagonal elements of the matrix under consideration can be expressed as a constant (one in this case) plus a random variable which is identical to the random variables of the off-diagonal terms, which is not the case. In spite of this, it was observed that this deviation from the assumptions in Baron et al. [2022] has not sizable impact on the eigenvalue spectra. When the elements of a random matrix are extracted from different distributions, the results in Poley et al. [2023] can be applied. By doing so, it can be seen that the diagonal deviation in the present case has an effect that rapidly vanishes with $N$, which explains the observed behavior. This is still, however, a matter of conjecture, because the potential correlations between elements with one index in common (type 2 correlations in Baron et al. [2022]) are not considered in Poley et al. [2023]. In this line, we think that the results presented in Baron et al. [2022] and Poley et al. [2023] are compatible at least for matrices with random components with zero mean, where according to Baron et al. [2022] type 2 correlations have no effect on the eigenvalue spectra.

Considering these all, let us assume that equation 4.5.2 is a good approximation of the region where the eigenvalues of $I + L^{-1}\Delta$ are distributed in the limit $N \to \infty$. Moreover, the contribution of $\langle {}_D\alpha_{\text{-};j,\text{-}} \rangle - \mathbb{E}\big[\langle {}_D\alpha_{\text{-};j,\text{-}} \rangle\big]$ to the variance of the terms $\Delta_{i,j}$ can be neglected if $N$ is big enough. Given this and the properties of variance and covariance:

$$\sigma^2_{\Delta_{i,j}} \sim \sum_{D=2}^{\infty} \binom{N-2}{D-2} \frac{\phi_D^2 C_D}{N^{2(D-2)}} \Big[1 - C_D\big(2P_D - 1\big)^2\Big] \tag{4.5.3}$$

which in the limit $N \to \infty$:

$$\sigma^2_{\Delta_{i,j}}\big(N \to \infty\big) = \sum_{D=2}^{\infty} \frac{\phi_D^2 C_D}{(D-2)!} \Big[1 - C_D\big(2P_D - 1\big)^2\Big] N^{2-D} \tag{4.5.4}$$

By combining equations 4.5.3 and 4.5.2 and substituting $\mathbb{E}\big[\langle_D\alpha_{-;-}\rangle\big] = \phi_D C_D\Big(2P_D - 1\Big)$ according to equations 4.1.9 and 4.1.5:

$$\frac{(x-1)^2}{(1+\rho(\Delta_{i,j},\Delta_{j,i}))^2} + \frac{y^2}{(1-\rho(\Delta_{i,j},\Delta_{j,i}))^2} = \frac{1}{(1+\beta)^2} \sum_{D=2}^{\infty} \binom{N-2}{D-2} \frac{\phi_D^2 C_D}{N^{2D-5}} \Big[1 - C_D\big(2P_D - 1\big)^2\Big]$$
$$\text{with } \beta = 1 + \sum_{D=2}^{\infty} \binom{N-2}{D-2} / N^{D-2} \phi_D C_D\big(2P_D - 1\big) \tag{4.5.5}$$

which in the limit $N \to \infty$:

$$\frac{(x-1)^2}{(1+\rho(\Delta_{i,j},\Delta_{j,i}))^2} + \frac{y^2}{(1-\rho(\Delta_{i,j},\Delta_{j,i}))^2} = \frac{1}{(1+\beta)^2} \sum_{D=2}^{\infty} \frac{\phi_D^2 C_D}{(D-2)!} \Big[1 - C_D\big(2P_D - 1\big)^2\Big] N^{3-D}$$
$$\text{with } \beta = 1 + \sum_{D=2}^{\infty} \frac{\phi_D C_D\big(2P_D - 1\big)}{(D-2)!} \tag{4.5.6}$$

Under the same approaches as those used for calculating $\sigma^2_{\Delta_{i,j}}$ and considering equation 4.1.14, $\rho(\Delta_{i,j},\Delta_{j,i})$ can be calculated from the network properties:

$$\rho(\Delta_{i,j},\Delta_{j,i}) \sim \frac{\sum_{D=2}^{\infty} \binom{N-2}{D-2} \frac{\phi_D^2 C_D}{N^{2(D-2)}} \bigg(\Big[2S_{_D} - 1 - C_D\big(2P_D - 1\big)^2\Big] - A_D\Big[2S_{_D} - 1\Big]\bigg)}{\sum_{D=2}^{\infty} \binom{N-2}{D-2} \frac{\phi_D^2 C_D}{N^{2(D-2)}} \Big[1 - C_D\big(2P_D - 1\big)^2\Big]} \tag{4.5.7}$$

which in the limit $N \to \infty$:

$$\rho(\Delta_{i,j},\Delta_{j,i}) = \frac{\sum_{D=2}^{\infty} \frac{\phi_D^2 C_D}{(D-2)!} \bigg(\Big[2S_{_D} - 1 - C_D\big(2P_D - 1\big)^2\Big] - A_D\Big[2S_{_D} - 1\Big]\bigg) N^{2-D}}{\sum_{D=2}^{\infty} \frac{\phi_D^2 C_D}{(D-2)!} \Big[1 - C_D\big(2P_D - 1\big)^2\Big] N^{2-D}} \tag{4.5.8}$$

#### 4.6 Upper bound for $\max_{x,y}\left[\frac{\left|\sum^{\binom{N}{D-2}} x_{k_1}...x_{k_{D-2}}-y_{k_1}...y_{k_{D-2}}\right|}{|x-y|_\infty}\right]$ and $\max_{i,x,y}\left[\frac{\left|\sum_{k_l\neq i}^{\binom{N-1}{D-2}} x_{k_1}...x_{k_{D-2}}-y_{k_1}...y_{k_{D-2}}\right|}{|x-y|_\infty}\right]$:

First of all, it is important to note that, because the objective is to find solutions in the ball $x \in X$, $|x-x_0|_\infty < 1/N$, with $x_0$ the vector with all components equal to $1/N$ (*i.e.* the homogeneous case), the vectors $x, y$ considered have their components bounded between 0 and $2/N$.

Let us start by bounding $\max_{x,y}\left[\frac{\left|\sum^{\binom{N}{D-2}} x_{k_1}...x_{k_{D-2}}-y_{k_1}...y_{k_{D-2}}\right|}{|x-y|_\infty}\right]$. It is clear that this maximum is zero for $D \leq 3$, so let us assume $D > 3$. By the mean value theorem in several variables:

$$\sum^{\binom{N}{D-2}} x_{k_1}...x_{k_{D-2}} - y_{k_1}...y_{k_{D-2}} = \nabla_{S_{D-2}}\big(cx + (1-c)y\big)\cdot(x-y) \tag{4.6.1}$$

with $c \in [0,1]$ and $\nabla_{S_{D-2}}$ the vector of partial derivatives of the function $\sum^{\binom{N}{D-2}} x_{k_1}...x_{k_{D-2}}$:

$$_i\nabla_{S_{D-2}}(z) = \sum_{k_l\neq i}^{\binom{N-1}{D-3}} z_{k_1}z_{k_2}...z_{k_{D-3}} \tag{4.6.2}$$

where $_i\nabla_{S_{D-2}}(z)$ indicates the $i^{th}$ component of $\nabla_{S_{D-2}}$ evaluated at vector $z$. Considering this expression, taking absolute values in both sides of equation 4.6.1 and operating:

$$\begin{aligned}\Big|\sum^{\binom{N}{D-2}} x_{k_1}...x_{k_{D-2}} - y_{k_1}...y_{k_{D-2}}\Big| &= \big|\nabla_{S_{D-2}}(cx+(1-c)y)\cdot(x-y)\big| \leq \max_z \big|\nabla_{S_{D-2}}(z)\cdot(x-y)\big| = \\ = \max_z\Big|\sum_i^N (y_i - x_i)\sum_{k_l\neq i}^{\binom{N-1}{D-3}} z_{k_1}z_{k_2}...z_{k_{D-3}}\Big| &= |x-y|_\infty \max_z\Big|\sum_i^N \delta_i \sum_{k_l\neq i}^{\binom{N-1}{D-3}} z_{k_1}z_{k_2}...z_{k_{D-3}}\Big| = |x-y|_\infty \max_z G_D\end{aligned} \tag{4.6.3}$$

where $\delta_i|x-y|_\infty = (y_i - x_i)$ which implies $\delta_i \in [-1,1]$. Note that, because $\sum_i^N x_i = 1$ by definition, them $\sum_i^N \delta_i = 0$. It is clear from inequality 4.6.3 that the maximum of $G_D$ is an upper bound of $\max_{x,y}\left[\frac{\sum^{\binom{N}{D-2}} |x_{k_1}...x_{k_{D-2}}-y_{k_1}...y_{k_{D-2}}|}{|x-y|_\infty}\right]$. The value of $\max G_D(z)$ can be bounded by induction:

$$\begin{aligned}G_{D+1} = \Big|\sum_i^N \delta_i \sum_{k_l\neq i}^{\binom{N-1}{D-2}} z_{k_1}z_{k_2}...z_{k_{D-2}}\Big| &= \frac{1}{D-2}\Big|\sum_i^N \delta_i \sum_{k_l\neq i}^{\binom{N-1}{D-3}} z_{k_1}z_{k_2}...z_{k_{D-3}}(1 - z_{k_1} - z_{k_2} - ... - z_{k_{D-3}})\Big| \leq \\ \leq \frac{1}{D-2}\Big|\sum_i^N \delta_i \sum_{k_l\neq i}^{\binom{N-1}{D-3}} z_{k_1}z_{k_2}...z_{k_{D-3}}\Big| &= \frac{1}{D-2}G_D\end{aligned} \tag{4.6.4}$$

But $G_4 = \left|\sum_i^N \delta_i \sum_{k\neq i}^{N-1} z_k\right| = \left|\sum_i^N \delta_i(1-x_i)\right| \leq 1$, the equality attained by making half of the pairs $(\delta_i, x_i)$ equal to $(1,0)$ and the other half equal to $(-1, 2/N)$ (if $N$ is odd the remaining $\delta_i$ takes the zero value for attaining the equality while keeping the constraints). Knowing the value of $\max_z G_4(z)$ and the recurrence displayed in 4.6.4 it can be concluded that:

$$\frac{\left|\sum^{\binom{N}{D-2}} x_{k_1}...x_{k_{D-2}} - y_{k_1}...y_{k_{D-2}}\right|}{|x-y|_\infty} \leq \max_z G_D \leq \frac{1}{(D-3)!} \tag{4.6.5}$$

The value of $\max_{i,x,y}\left[\frac{\left|\sum_{k_l\neq i}^{\binom{N-1}{D-2}} x_{k_1}...x_{k_{D-2}} - y_{k_1}...y_{k_{D-2}}\right|}{|x-y|_\infty}\right]$ can be bounded in a completely analogous way, but with slight differences. The first one is that, in this case, the value of the maximum is zero for $D=2$ but one for $D=3$. Now, let us focus in cases $D>3$. By the mean value theorem in several variables:

$$\sum_{k_l\neq i}^{\binom{N-1}{D-2}} x_{k_1}...x_{k_{D-2}} - y_{k_1}...y_{k_{D-2}} = \nabla_{iS_{D-2}}\big(cx+(1-c)y\big)\cdot(x-y) \tag{4.6.6}$$

with $c\in[0,1]$ and $\nabla_{iS_{D-2}}$ the vector of partial derivatives of the function $\sum_{k_l\neq i}^{\binom{N-1}{D-2}} x_{k_1}...x_{k_{D-2}}$:

$${}_k\nabla_{iS_{D-2}}(z) = \sum_{k_l\neq i,k}^{\binom{N-2}{D-3}} z_{k_1}z_{k_2}...z_{k_{D-3}} \tag{4.6.7}$$

where ${}_k\nabla_{iS_{D-2}}(z)$ indicates the $k^{th}$ component of $\nabla_{iS_{D-2}}$ evaluated at vector $z$. Considering this expression, taking absolute values in both sides of equation 4.6.6 and operating:

$$\left|\sum_{k_l\neq i}^{\binom{N-1}{D-2}} x_{k_1}...x_{k_{D-2}} - y_{k_1}...y_{k_{D-2}}\right| = \left|\nabla_{iS_{D-2}}\big(cx+(1-c)y\big)\cdot(x-y)\right| \leq \max_z\left|\nabla_{iS_{D-2}}(z)\cdot(x-y)\right| =$$
$$= \max_z\left|\sum_{k\neq i}^{N-1}(y_k-x_k)\sum_{k_l\neq i,k}^{\binom{N-1}{D-3}} z_{k_1}z_{k_2}...z_{k_{D-3}}\right| = |x-y|_\infty \max_z\left|\sum_{k\neq i}^{N-1}\delta_k\sum_{k_l\neq i,k}^{\binom{N-2}{D-3}} z_{k_1}z_{k_2}...z_{k_{D-3}}\right| = |x-y|_\infty \max_z {}_iG_D \tag{4.6.8}$$

where, again, $\delta_k|x-y|_\infty = (y_k - x_k)$. Given $i$ is fixed, the maximum of ${}_iG_D$ is an upper bound of $\max_{x,y}\left[\frac{\sum_{k_l\neq i}^{\binom{N-1}{D-2}} |x_{k_1}...x_{k_{D-2}} - y_{k_1}...y_{k_{D-2}}|}{|x-y|_\infty}\right]$. The value of $\max {}_iG_D(z)$ can be bounded by induction as before:

$${}_iG_{D+1} = \left|\sum_{k\neq i}^{N-1}\delta_k\sum_{k_l\neq i,k}^{\binom{N-2}{D-2}} z_{k_1}z_{k_2}...z_{k_{D-2}}\right| = \frac{1}{D-2}\left|\sum_{k\neq i}^{N-1}\delta_k\sum_{k_l\neq i,k}^{\binom{N-2}{D-3}} z_{k_1}z_{k_2}...z_{k_{D-3}}\big(1-x_i-z_{k_1}-z_{k_2}-...-z_{k_{D-3}}\big)\right| \leq$$
$$\leq \frac{1}{D-2}\left|\sum_{k\neq i}^{N-1}\delta_k\sum_{k_l\neq i,k}^{\binom{N-2}{D-3}} z_{k_1}z_{k_2}...z_{k_{D-3}}\right| = \frac{1}{D-2}{}_iG_D \tag{4.6.9}$$

But ${}_iG_{{}_4} = \Big| \sum_{k\neq i}^{N-1} \delta_k \sum_{j\neq i,k}^{N-2} z_j \Big| = \Big| \sum_{k\neq i}^{N-1} \delta_k \big(1 - x_i - x_k\big) \Big| = \Big| - \delta_i \big(1 - x_i\big) + \sum_{k\neq i}^{N-1} \delta_k x_k \Big| < 2$. The equality would be attained by making $x_i = 0$, $\delta_i = -1$, $N/2 - 1$ of the pairs $\big(\delta_k, x_k\big)$ equal to $\big( -1, \frac{2}{N-2}\big)$ and the remaining $N/2$ pairs equal to $\big(1, 0\big)$ if $N$ is even; or by making $x_i = 0$, $\delta_i = -1$, $(N-1)/2-1$ of the pairs $\big(\delta_k, x_k\big)$ equal to $\big(-1, \frac{2}{N-3}\big)$,$(N-1)/2$ pairs equal to $\big(1, 0\big)$ and the remaining pair equal to $\big(0, 0\big)$ if $N$ is odd. Note that the equality cannot be achieved because $\frac{2}{N-2}$ and $\frac{2}{N-3}$ are greater than $\frac{2}{N}$, the maximum value of $x_i$ allowed. Knowing this upper value of $\max_z {}_iG_{{}_4}(z)$ and the recurrence displayed in 4.6.4 it can be concluded that:

$$\frac{\Big| \sum_{k_l\neq i}^{\binom{N-1}{D-2}} x_{k_1}...x_{k_{D-2}} - y_{k_1}...y_{k_{D-2}} \Big|}{|x-y|_\infty} \leq \max_z {}_iG_{{}_D} \leq \frac{2}{(D-3)!} \tag{4.6.10}$$

#### 5 Supplementary Figures.

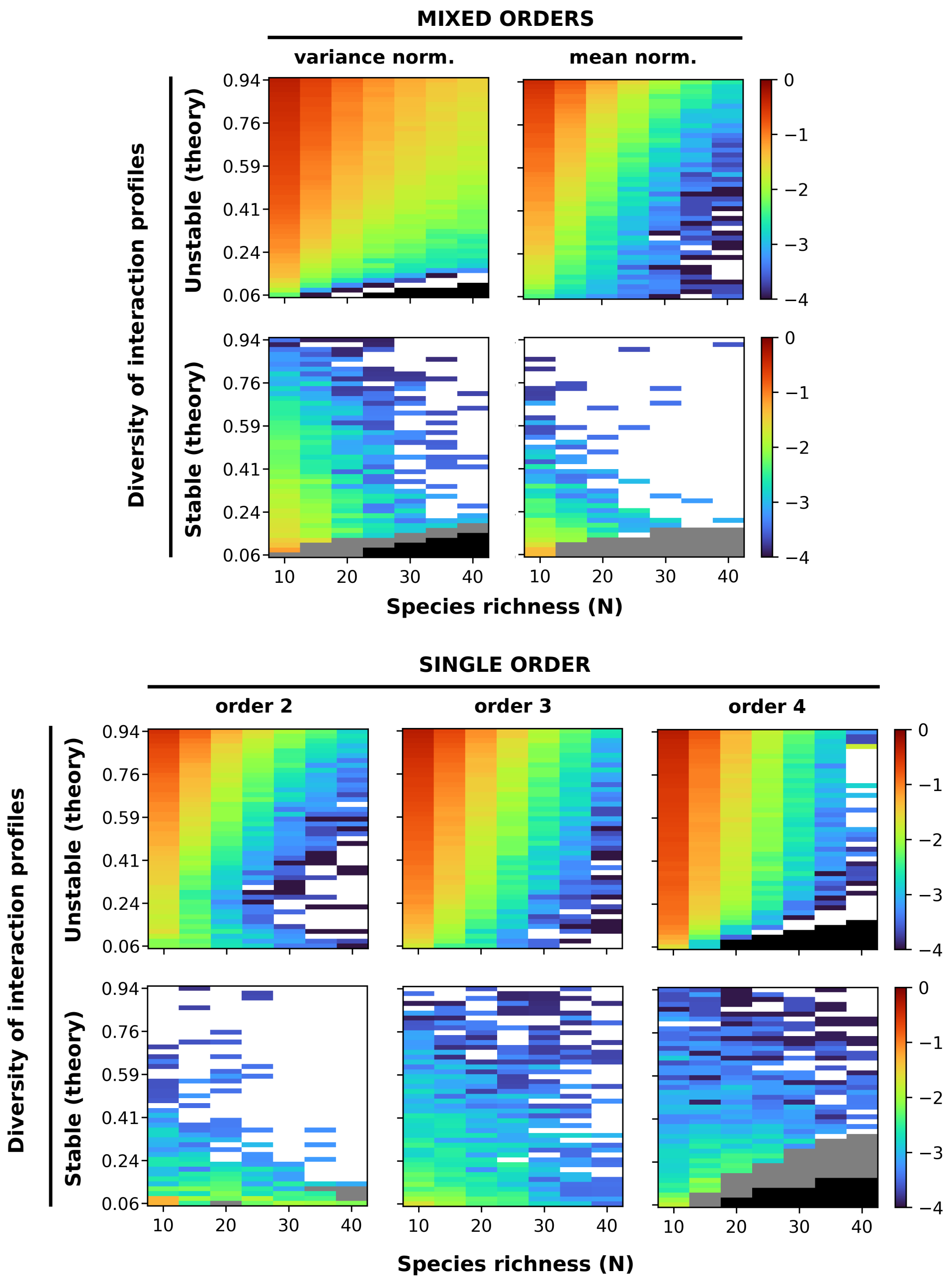

Figure 1: Validity of the approximate stability criterion (eqn. 2 in the main text and eqns. 3.1.7 in the Supplementary Information). The color maps represent the logarithm (base 10) of the fraction of simulations that are predicted to be stable (unstable) based on the stability criterion but are actually unstable (stable), for different interaction orders, species richness ($N$) and diversity of interaction profiles. The upper plots correspond to a mixture of interactions of order 2, 3, and 4, with interaction strengths scaled to obtain balanced means (mean normalization) or variances (variance normalization, see Supplementary Information section 3). The colder the color, the better the approximation; white color indicates no errors. A total of 9604 simulations were performed for each tile, which allows estimation of proportions with a 95% confidence interval narrower than ±0.01 under the normal approximation. Black regions indicate unfeasible parameter combinations. Gray regions correspond to parameter combinations in which most simulated networks lacked a stationary state with full coexistence (stable or unstable), precluding any further assessment of stability.

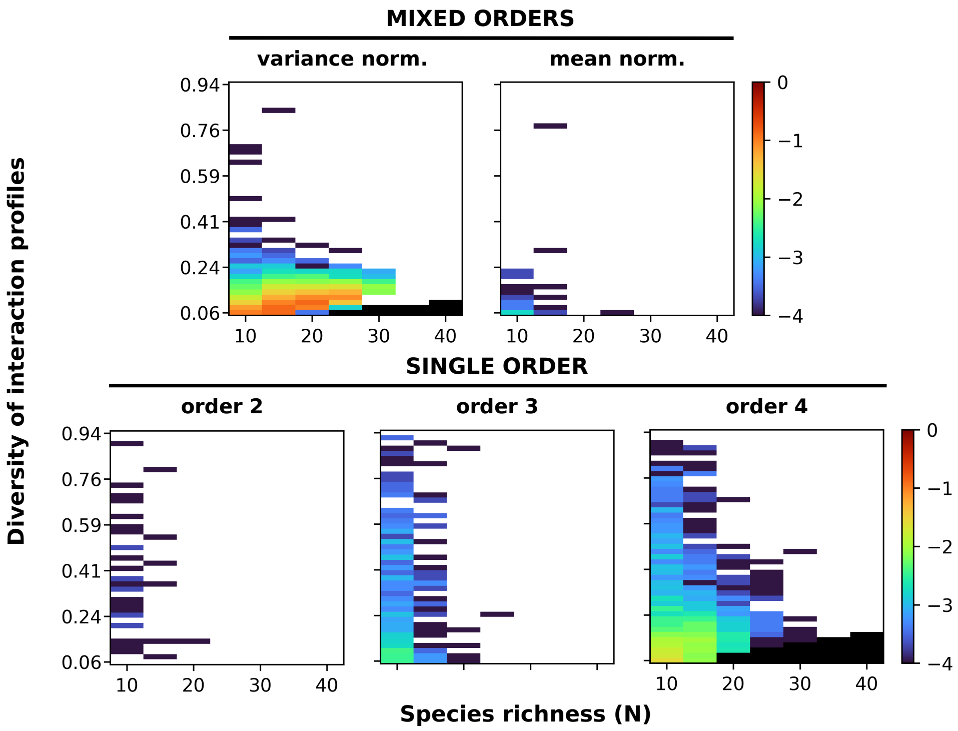

Figure 2: Frequency of potential non-fixed-point attractors. The color maps represent the logarithm (base 10) of the fraction of simulations that fulfill the entropy criterion but not the derivative criterion (see Methods: Numerical simulations of community dynamics). The colder the color, the less likely to find these attractors; white color corresponds to zero cases. A total of 9604 simulations were performed for each tile, which allows estimation of proportions with a 95% confidence interval narrower than $\pm 0.01$ under the normal approximation. Black regions indicate unfeasible parameter combinations.

### References

Ioannis K. Argyros and Hongmin Ren. A simplified proof of the kantorovich theorem for solving equations using telescopic series. *J. Numer. Anal. Approx. Theory*, 44(2):146–153, 2015. doi: 10.33993/jnaat442-1084.

Joseph W. Baron, Thomas Jun Jewell, Christopher Ryder, and Tobias Galla. Eigenvalues of random matrices with generalized correlations: A path integral approach. *Phys. Rev. Lett.*, 128(120601): 1–6, 2022. doi: 10.1103/PhysRevLett.128.120601.

Jiu Ding and Aihui Zhou. Eigenvalues of rank-one updated matrices with some applications. *Appl. Math. Lett.*, 20(12):1223–1226, 2007. doi: 10.1016/j.aml.2006.11.016.

J.G. Expectation of maximum of n i.i.d random variables. Mathematics Stack Exchange, 2019. URL https://math.stackexchange.com/q/3305183(version:2019-07-27).

Grégoire Lecerf and Joelle Saadé. A short survey on kantorovich-like theorems for newton's method. *ACM Commun. Comput. Algebra*, 50(1):1–11, 2015. doi: 10.1145/2930964.2930965.

Lyle Poley, Tobias Galla, and Joseph W. Baron. Eigenvalue spectra of finely structured random matrices. *arXiv*, (2311.02006), 2023. doi: 10.48550/arXiv.2311.02006.

J.M. Varah. A lower bound for the smallest singular value of a matrix. *Linear Algebra Appl.*, 11(1): 3–5, 1975. doi: 10.1016/0024-3795(75)90112-3.
